# Discovery of Potent Degraders of the Dengue Virus Envelope Protein

**DOI:** 10.1101/2024.06.01.596987

**Authors:** Zhengnian Li, Han-Yuan Liu, Zhixiang He, Antara Chakravarty, Ryan P. Golden, Zixuan Jiang, Inchul You, Hong Yue, Katherine A. Donovan, Guangyan Du, Jianwei Che, Jason Tse, Isaac Che, Wenchao Lu, Eric S. Fischer, Tinghu Zhang, Nathanael S. Gray, Priscilla L. Yang

## Abstract

Targeted protein degradation has been widely adopted as a new approach to eliminate both established and previously recalcitrant therapeutic targets. Here we report the development of small molecule degraders of the envelope (E) protein of dengue virus. We developed two classes of bivalent E-degraders, linking two previously reported E-binding small molecules, GNF-2 and CVM-2-12-2, to a glutarimide-based recruiter of the CRL4^CRBN^ ligase to effect proteosome-mediated degradation of the E protein. ZXH-2-107 (based on GNF-2) is an E degrader with ABL inhibition while ZXH-8-004 (based on CVM-2-12-2) is a selective and potent E-degrader. These two compounds provide proof-of-concept that difficult-to-drug targets such as a viral envelope protein can be effectively eliminated using a bivalent degrader and provide starting points for the future development of a new class antiviral drugs.

## Introduction

Dengue is a mosquito-borne tropical disease caused by dengue virus (DENV) infection. Dengue disease has affected more than 100 countries in tropical and subtropical regions, putting approximately 3.6 billion people at risk.^1, 2^ It was estimated that approximately 390 million people are infected per year, with an estimated 96 million people experiencing severe disease.^3^ Dengue virus belongs to the genus Flavivirus of the Flaviviridae family, which includes several other human pathogens including Zika, yellow fever, West Nile and Japanese encephalitis viruses (ZIKV, YFV, WNV, and JEV, respectively).^4, 5^

Dengue virus strains are classified into four related serotypes of virus (DENV1-4).^6^ The genome of DENV encodes three structural proteins (the envelope (E) protein, precursor membrane/ membrane (prM) / (M) protein, and capsid (C) protein), which are essential components of the viral particle, and seven non-structural (NS) proteins (NS1, NS2A, NS2B, NS3, NS4A, NS4B and NS5) that are responsible for replication of the viral genome and evasion of the host immune system.^7^

To date, there are still no approved antiviral agents to prevent or treat DENV infection. Although the first dengue vaccine, Dengvaxia, developed by Sanofi Pasteur was licensed in December 2015, follow-up studies demonstrated an increased risk of severe dengue and increased risk of hospitalization in seronegative individuals after Dengvaxia vaccination.^8^ Therefore, there remains an urgent need for new vaccines and small molecule anti-viral agents.

The envelope (E) protein covers the surface of the DENV virion as 90 prefusion dimers atop the virus’s lipid membrane. The E protein is comprised of three distinct structural domains, EDI, EDII, and EDIII.^9, 10^ During viral entry, the initial attachment step is mediated by binding of EDIII on the virion to host factors on the plasma membrane surface. Following endocytosis of the virion, acidification of the endosomal compartment leads to structural changes in E that are coupled to fusion of the viral and endosomal membranes, culminating in the formation of a pore that allows the viral genome to escape to the cytosol. The DENV2 E protein was revealed to have a hydrophobic, ligand-binding pocket located in the hinge region between EDI and EDII when a molecule of octyl-β-D-glucoside (β-OG) included in the crystallization solution co-crystallized with the protein.^11^ Given the essential role of E in mediating both the attachment and fusion steps of virus entry, targeting the β-OG pocket of the E protein with small molecules was proposed as a potential strategy for the development of a new class of anti-viral agents.^12-14^ Previously, our group reported that the allosteric ABL kinase inhibitor (GNF-2), which targets the myristate-binding pocket in BCR-ABL, interferes with E-mediated membrane fusion during viral entry, with both pharmacological activities contributing to antiviral activity.^15^ Photoaffinity crosslinking studies and the identification of resistance mutations in the β-OG pocket subsequently identified this site as the molecular target of these compounds.^16^ A subsequent medicinal chemistry effort aimed at reducing binding to BCR-ABL resulted in the development of CVM-2-12-2, a 2,4-diamino-substituted pyrimidine, and other analogs with increased activity against DENV2 entry compared to GNF-2.^15, 16^ Although GNF-2 and CVM-2-12-2 provide proof-of-concept for antiviral activity through small molecules targeting E, the antiviral potency of these molecules^15, 16^ is currently insufficient to meet the extremely high antiviral potency required for effective antiviral drugs.

Bifunctional degraders, also known as Proteolysis Targeting Chimeras (PROTACs), have recently emerged as a promising alternative targeting modality in drug discovery.^17-19^ PROTACs are comprised by two protein-binding ligands, one that recruits an E3 ligase and the other that binds a protein of interest (POI). Formation of the resulting ternary complex enables ubiquitination and subsequent proteasomal degradation of the POI. This strategy has been applied most frequently and successfully in oncology, with more than a dozen PROTAC drugs entering the clinical development stage by the end of 2021.^20^ In contrast, application of targeted protein degradation in the area of infectious disease is still an incipient field of research.^21^ Recently, our group developed the first small molecule anti-viral PROTACs by using telaprevir, an established direct-acting antiviral targeting the hepatitis C virus (HCV) NS3-4A protease, to develop bifunctional degraders.^22^ Subsequently, PROTAC molecules designed to target the influenza virus neuraminidase,^23^ hemagglutinin,^24^ PA endonuclease protein,^25^ and the SARS Cov-2 main protease (MPro),^26, 27^ have been reported. In addition, peptide-based PROTACs targeting the SARS CoV-2 spike protein,^28^ and hepatitis B virus X protein^29^ have been described. With only this limited set of examples, the repertoire of viral proteins susceptible to targeted protein degradation remains largely uncharacterized. Since viruses have evolved mechanisms to ensure robust expression of their genomes, it is unclear whether highly abundant viral proteins, such as the envelope protein and other structural proteins that are needed for production of progeny virions, can be depleted sufficiently to achieve significant antiviral activity. In addition, many viral proteins may not be susceptible due to their subcellular localization.

Here, we report the development of dengue E protein-directed PROTACs consisting of pyrimidine-derived E-inhibitors (GNF-2, CVM-2-12-2) conjugated to glutarimide-derived recruiters of the E3 CRL4^CRBN^ ubiquitin ligase. A structure-activity exploration of these two E-binders resulted in the discovery of ZXH-2-107 and ZXH-8-004 which achieve potent on-target degradation of the E protein. Both ZXH-2-107 and ZXH-8-004 exhibit antiviral activity against DENV2 in cell culture that is significantly higher than that of their respective parental inhibitors.

## Results and Discussion

### Development of GNF-2-based E protein degrader

To develop an E protein-targeted degrader, we started with modification of GNF-2 (Figure 1A), conjugating it to a CRBN-binding ligand derived from thalidomide. Our strategy was informed by computational docking of GNF-2 into the hydrophobic octyl-β-D-glucoside (β-OG)-binding pocket of E using Glide (Schrödinger suite) (Figure 1B). The model predicts that the 4-(trifluoromethoxy) phenyl moiety is deeply embedded in the hydrophobic pocket and that the -NH of GNF-2 forms a hydrogen bond with Thr48 within the pocket. In contrast, the amide group protrudes outside and is solvent-exposed, forming three hydrogen-bonding interactions with Gln200, Asp 203, and Ser274 near the top of the pocket. These modeling studies suggested that the amide bond is an appropriate site for the attachment of linkers.

**Figure 1.**
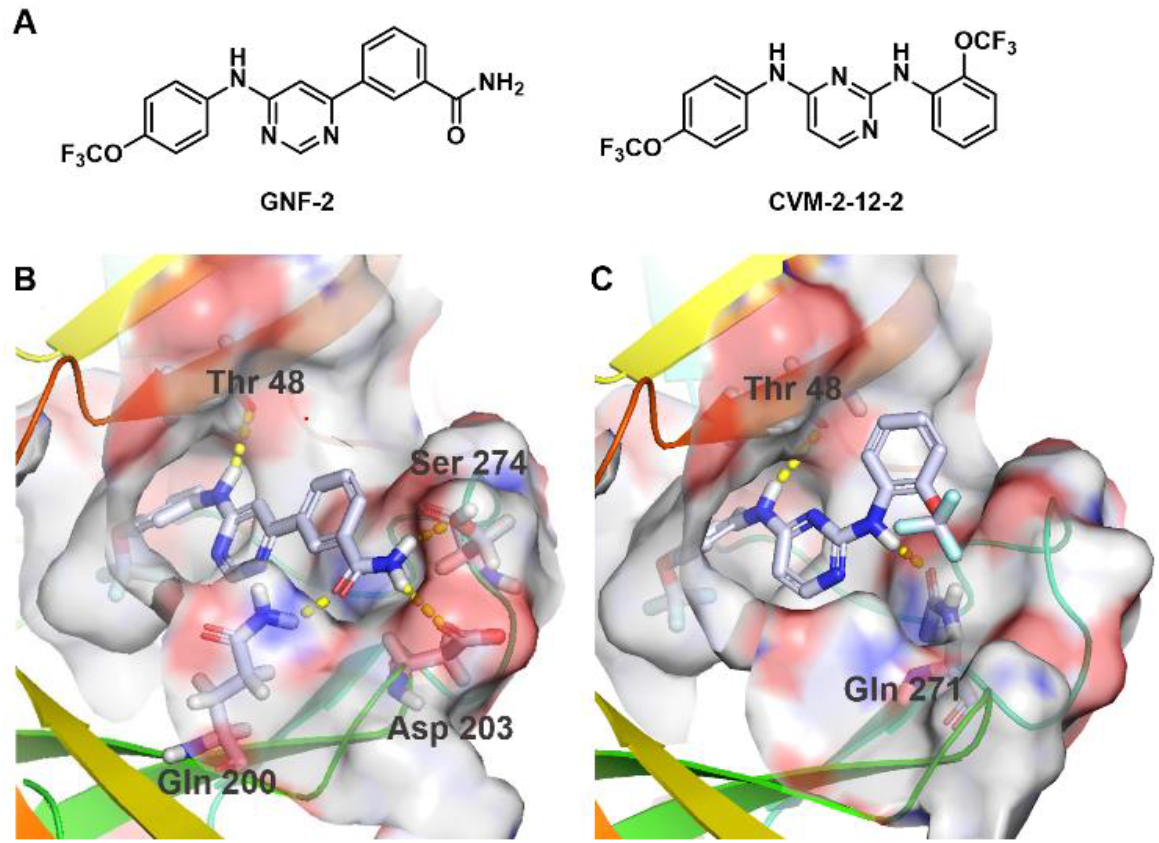
Design of PROTACs targeting the dengue E protein. A) Chemical structures of GNF-2 and CVM-2-12-2. B) Optimal docking poses of dengue E protein (PDB: 1OKE) with GNF-2. C) Optimal docking poses of dengue E protein (PDB: 1OKE) with CVM-2-12-2. The E protein is shown as a ribbon cartoon and the key residues forming hydrogen bonds are represented as sticks. Docking studies were performed with Schrödinger, and models were prepared with PyMOL.

Accordingly, we synthesized a series of candidate dengue E-targeting degrader molecules by conjugating GNF-2 through the solvent-exposed amide position. The E3 ligase ligands of CRBN were successfully used in our previous HCV NS3/4A protease PROTACs and were examined in this work.^22^ Compounds were synthesized by amidation from the 3-(pyrimidin-4-yl) benzoic acid, linked with either a polyethylene glycol (PEG) or an alkyl linker, and terminated at the 4-position or 5-position of thalidomide (Table 1). To assess whether these compounds induced degradation of dengue E in cells, we examined their effects on E abundance in DENV2-infected cells. For this, Huh 7.5 cells were infected at a multiplicity of infection (MOI) of 1 for one hour, and then incubated in the presence of the candidate degraders. Cell lysates and culture supernatants were harvested at 24 hours post-infection to allow examination of intracellular E abundance by immunoblot and quantification of infectious progeny virus, respectively (Figure 2A).

**Table 1.** The structures of GNF-2 based dengue E degraders.

| Compound | Linker | E3 recruiter |
| --- | --- | --- |
| ZXH-2-101 |  | B |
| ZXH-2-107<br>(GNF-2-deg) |  | A |
| ZXH-2-102 |  | B |
| ZXH-2-105 |  | A |
| ZNL-05-199 |  | B |
| ZXH-2-107-Neg<br>(GNF-2-deg-BUMP) |  | C |

**Figure 2.**
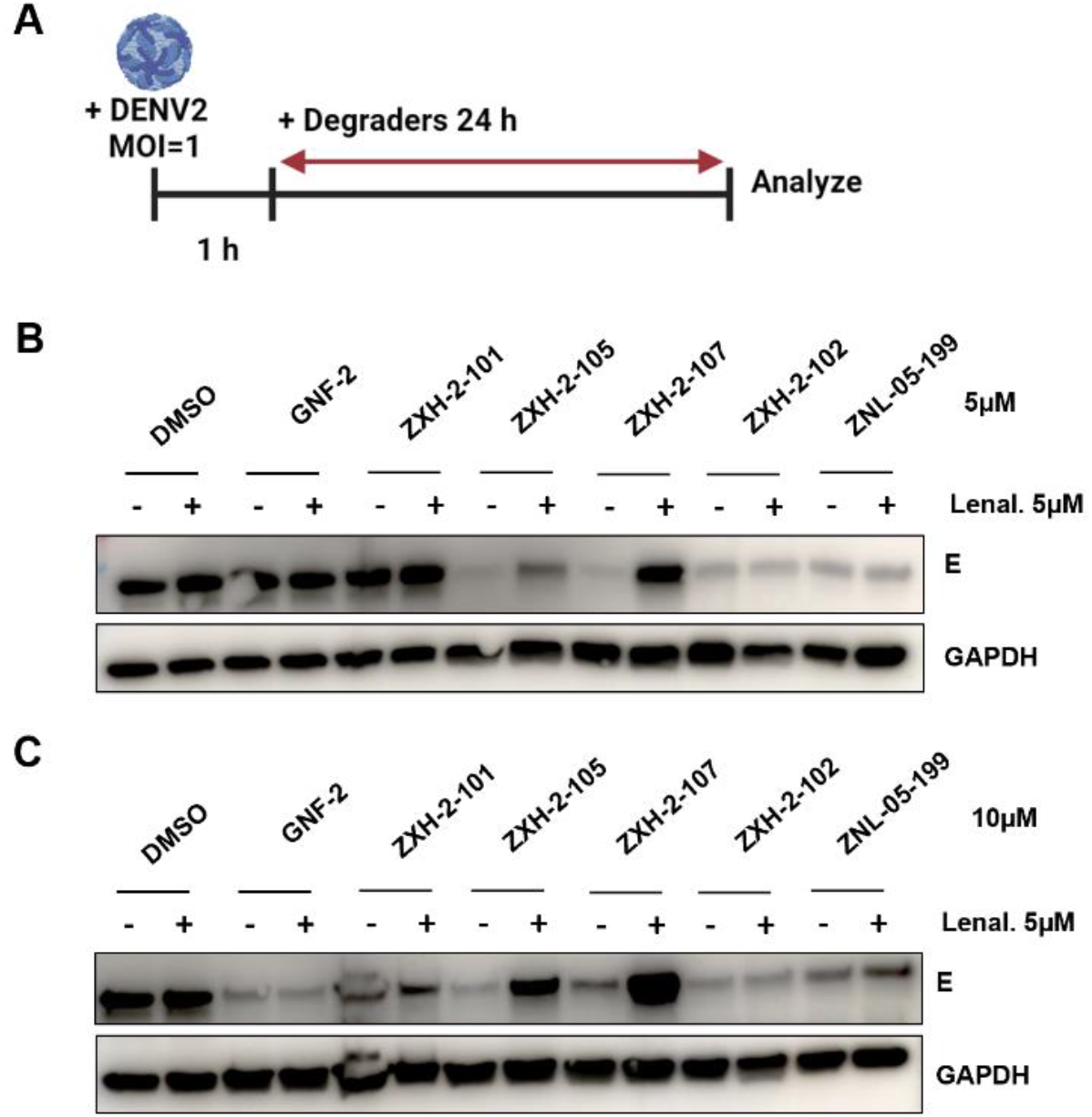
Screening of GNF-2 based degraders. A) Schematic representation of the viral infection assay utilized to evaluate E protein degradation and antiviral activity. Created on BioRender.com with permission. B) and C) Immunoblot analyses of dengue E abundance following treatment of DENV2-infected Huh7.5 cells with GNF-2-based degraders at 5 and 10 μM with and without pretreatment of 5 μM lenalidomide.

As shown in Figures 2B and 2C, several compounds caused reduction of E detected in infected cell lysates following a 24 h treatment of infected Huh 7.5 cells at 5 μM and 10 μM. This time point was chosen to allow correlation of degradation of E with antiviral activity, as 24 h represents the earliest time progeny virus – the product of the viral infectious cycle – is measurable under these experimental conditions. GNF-2 also caused a strong reduction in E protein levels at 10 μM because it inhibits E-mediated membrane fusion, thereby blocking viral entry and preventing expression of E and additionally exerts antiviral activity through inhibition of ABL kinases.^15^ To evaluate whether the candidate E PROTACs exert antiviral activity through a targeted protein degradation mechanism, we performed competitive assays in which treatment of DENV-infected Huh 7.5 cells was performed in the presence of an excess of the CRBN-binding ligand lenalidomide. Immunoblotting showed a large reduction of E protein in lysates from cells treated with ZXH-2-107 alone (Figure 2B & 2C); however, this depletion of E was blocked in the presence of lenalidomide, indicating that the engagement of CRBN is essential for depletion of E in the presence of ZXH-2-107, and suggesting that this compound causes CRBN-dependent depletion of E. Interestingly, ZXH-2-105 also induced strong reduction of E, however, this was only partially rescued in the presence of excess lenalidomide, which suggests that it has a different antiviral mechanism. Furthermore, ZXH-2-102 and ZNL-05-199 caused reductions in E both with and without lenalidomide pretreatment, indicating that these compounds also possess pharmacology that is independent of E3 ligase recruitment.

To characterize ZXH-2-107’s antiviral activity, we considered several potential antiviral mechanisms. Parental inhibitor GNF-2 and related compound GNF-5 are allosteric inhibitors of BCR-ABL and other ABL-family kinases that bind in the myristate-pocket^30^ and have been previously used to develop cIAP (cellular inhibitor of apoptosis protein)-derived PROTACs that induce degradation of BCR-ABL.^31, 32^ As mentioned previously, parental inhibitor GNF-2 itself has a dual mechanism of antiviral activity action, binding to E on incoming virions and inhibiting E-mediated membrane fusion during viral entry while also inhibiting a post-entry step in viral replication through its inhibition of cellular ABL kinases.^15^ Therefore, ZXH-2-107 could exert antiviral effects through inhibition or targeted degradation of ABL-family kinases and/or inhibition or targeted degradation of dengue E.

To determine whether ZXH-2-107 exerts antiviral activity through targeted degradation of ABL-family kinases, we tested whether ZXH-2-107 induces ABL degradation in the Huh 7.5 cells in a concentration-dependent manner. Immunoblot analysis revealed no reduction of ABL protein levels within 8 hours of treatment with ZXH-2-107 even at a concentration of 10 μM (Figure S1). We further examined ZXH-2-107 for targeted degradation of ABL in K562 cells, which are chronic myelogenous leukemia (CML)-derived cells that are dependent upon high expression of ABL-family kinases and thus highly sensitive to ABL depletion. We did not observe depletion of ABL in the presence of up to 10 μM ZXH-2-107 treatment (Figure S2). These data suggest that ZXH-2-107 is not an ABL degrader.

To evaluate whether the inhibition of ABL kinases contributes to ZXH-2-107’s antiviral activity, we conducted antiproliferative assays in K562 cells over a 72-hour period. GNF-2 exhibited an IC_50_ of 0.53 μM in these experiments, whereas ZXH-2-107 exhibited significantly less potent activity with an IC_50_ of 6.6 μM (Figure S3), which is correlated with its decreased cellular permeability (Table S1). ZXH-2-107 demonstrated mild inhibitory effects on K562 cell proliferation through ABL inhibition. These experiments suggest that ZXH-2-107’s antiviral derives from targeted degradation of the E and also possibly through the inhibition of ABL-family kinases.

### Development and optimization of a potent E protein degrader

To enable a more straightforward evaluation of the antiviral potential of targeted degradation of E in the absence of ABL inhibition exhibited by ZXH-2-107, we sought to develop an independent E degrader series using an E inhibitor devoid of ABL-binding activity. For this we chose the 2,4-diamino pyrimidine scaffold CVM-2-12-2. Docking of CVM-2-12-2 suggested that CVM-2-12-2 engages Thr48, with a binding pose similar to that of GNF-2, while forming a key hydrogen-bonding interaction with Gln271. The model predicts that the *meta*-position of the trifluoromethoxy functionality in CVM-2-12-2 extends out of the pocket and is solvent-exposed (Figure 1C).

To make CVM-2-12-2-based degraders, we introduced a carboxylic acid at the *meta*-position of the trifluoromethoxy group and conjugated it with linkers via amidation reaction, using thalidomide as the CRBN-recruiting moiety. All CVM-2-12-2-based degrader molecules were evaluated with and without lenalidomide treatment in DENV2-infected Huh 7.5 cells (MOI 1) for 24 hours (Figure 3A & 3B). ZXH-8-004, which contains a PEG2 linker, demonstrated the most potent depletion of dengue E after 24 h treatment at both 5 μM and 10 μM. Similar to GNF-2, CVM-2-12-2 caused a reduction of E at 5 and 10 μM concentrations due the antiviral activity it exerts on E-mediated fusion during viral entry, which is notably not CRBN-dependent. In contrast, depletion of E by candidate PROTACs is CRBN-dependent, as demonstrated by the full rescue of E degradation in the presence of excess lenalidomide. ZXH-08-001, which differs from ZXH-08-004 only in the linkage site of the thalidomide moiety, was also observed to cause weak depletion of E at 10 μM. In our study, compounds that harbor the 4-substituted thalidomide (ZXH-8-004) induced greater targeted degradation of E than those with the thalidomide at the 5-position (ZXH-8-001), presumably because the location of the E3-targeting moiety influences the conformation of the ternary complex. ZXH-8-004 did not induce depletion of ABL as determined by immunoblot analysis in both DENV2-infected Huh 7.5 cells (Figure S1) and K562 cells (Figure S2). Consistent with the lack of interaction of CVM-2-12-2 with ABL family kinases, ZXH-8-004 showed very weak antiproliferative activities K562 cells (Figure S3).

**Figure 3.**
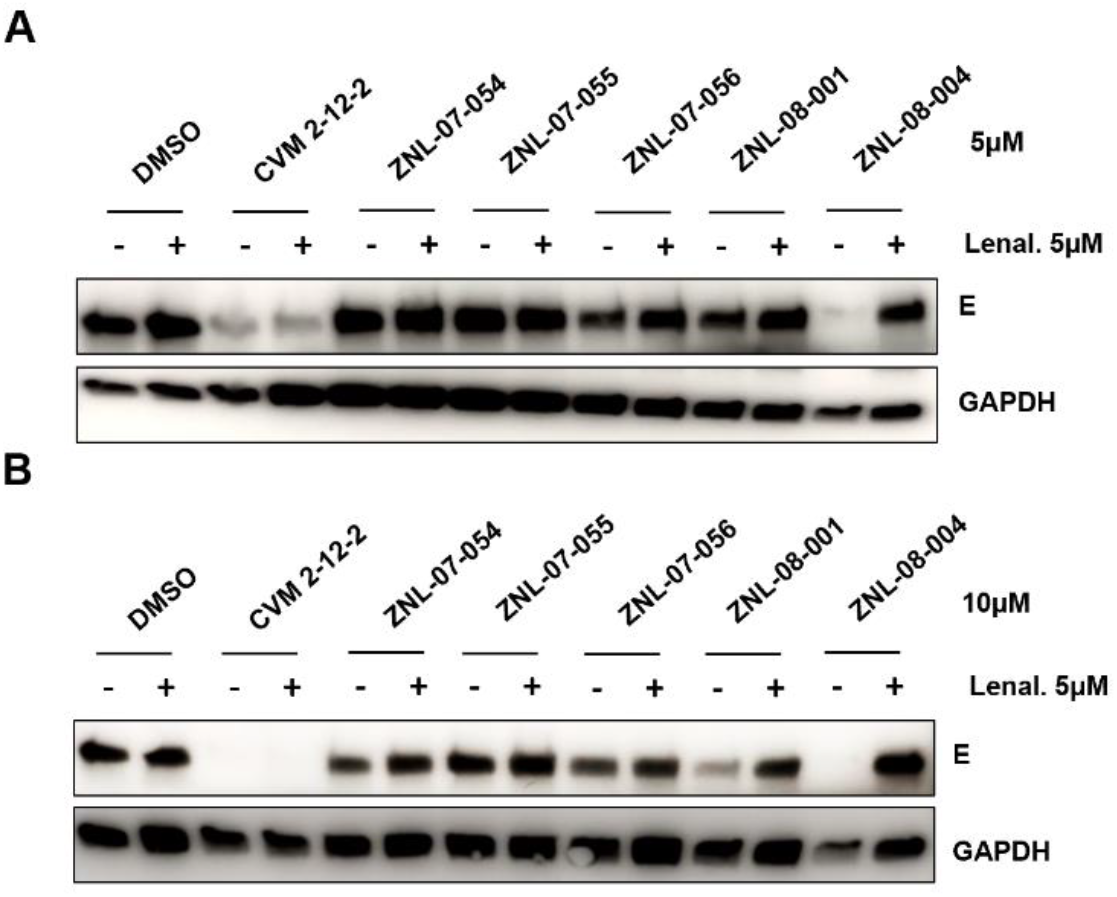
Screening of CVM-2-12-2 based degraders. A) Schematic representation of the viral infection assay utilized to evaluate E protein degradation and antiviral activity. B) and C) Immunoblot analyses of dengue E abundance following treatment of DENV2-infected Huh7.5 cells with CVM-2-12-2-based degraders at 5 and 10 μM with and without pretreatment of 5 μM lenalidomide.

Having successfully identified two chemically distinct scaffolds that could be used as targeting ligands for the development of bivalent degraders of dengue E, we performed additional experiments to characterize the mechanism of action of E degraders ZXH-2-107 and ZXH-8-004. For this, we synthesized corresponding negative control compounds (ZXH-2-107-Neg and ZXH-8-004-Neg) in which the glutarimide ring on thalidomide is replaced with a δ-lactam moiety, a modification that abrogates binding to CRBN.^33^ These negative control compounds provided us with the ability to compare ZXH-2-107 and ZXH-8-004 and to distinguish between inhibition- and degradation-dependent pharmacology (Table 1 and Table 2).

**Table 2.** The structures of CVM-2-12-2 based dengue E degraders.

| Compound | Linker | E3 recruiter |
| --- | --- | --- |
| ZNL-07-054 |  | B |
| ZNL-07-055 |  | B |
| ZNL-07-056 |  | B |
| ZXH-8-001 |  | A |
| ZXH-8-004<br>(2-12-2-deg) |  | B |
| ZXH-8-004-Neg<br>(2-12-2-deg-BUMP) |  | D |

Figure 4. A) Cellular CRBN engagement assay of ZXH-2-107 and ZXH-8-004 at 10 μM. B) E protein cellular thermal shift assay (CETSA) for E inhibitors and degraders. The quantitative analysis of immunoblot of sample from 59.7 ºC is performed with normalization of GAPDH.

### Engagement of E degraders with both CRBN and E protein

To measure the CRBN engagement for the two E degraders and negative controls in cells, we used a previously described competitive intracellular CRBN engagement assay.^34^ In this assay we measured the ability of degrader molecules to protect BRD4_BD2_ from degradation by dBET6, a pan-BET bromodomain degrader. The assay provides a measure of the cell penetrance of degraders as well as a measure of the compounds’ ability to engage CRBN intracellularly. As expected, neither ZXH-2-107-Neg nor ZXH-8-004-Neg exhibited evidence of CRBN engagement (Figure 4A). In contrast, both ZXH-2-107 and ZXH-8-004 engage CRBN intracellularly but not as potently as lenalidomide, most likely due to their increased molecular weights and reduced cell penetrance. Collectively, these data support that ZXH-2-107 and ZXH-8-004 are cell permeable and able to engage the CRL4^CRBN^ E3 ligase.

**Figure 4.**
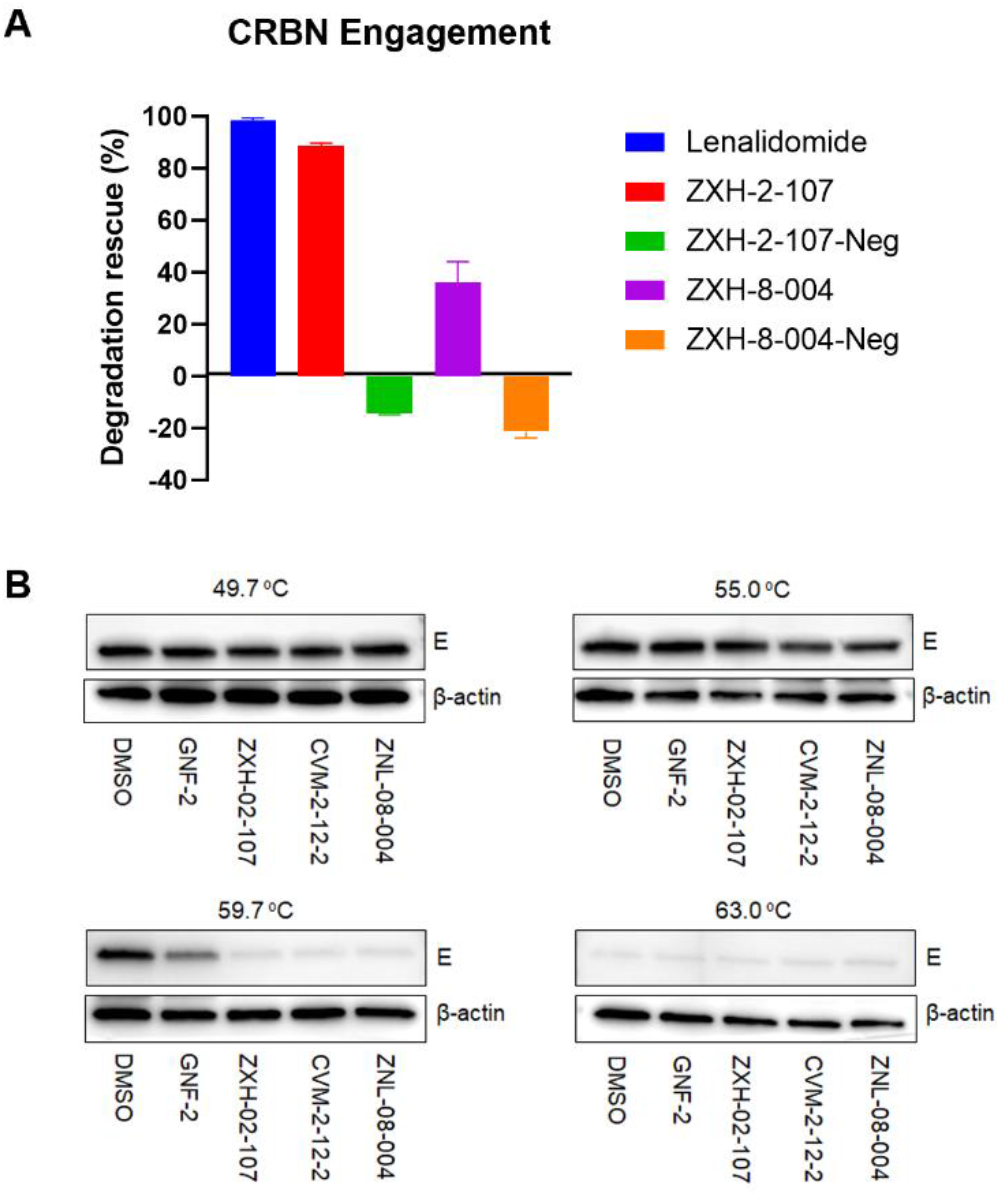
A) Cellular CRBN engagement assay of ZXH-2-107 and ZXH-8-004 at 10 μM. B) E protein cellular thermal shift assay (CETSA) for E inhibitors and degraders. The quantitative analysis of immunoblot of sample from 59.7 ºC is performed with normalization of GAPDH.

To assess whether ZXH-2-107 and ZXH-8-004 induced targeted degradation of dengue E by engagement of E in the cellular context, we performed cellular thermal shift assays (CETSA) in DENV2-infected Huh 7.5 cells.^35, 36^ We observed thermal-destabilized bands in the area of 53 kDa, the anticipated size of monomeric E, in samples treated with 10 μM of GNF-2, ZXH-2-107, CVM-2-12-2, and ZXH-8-004 and when heated at 49.7ºC, 55ºC, 59.7ºC, and 63 ºC. Immunoblot analysis showed significant thermal destabilization of E upon treatment with both the inhibitors and the degraders (Figure 4B). These results demonstrate that ZXH-2-107 and ZXH-8-004 both engage E directly in cells.

### ZXH-8-004 demonstrates potent antiviral activity as a CRBN-mediated E Degrader

We next examined the concentration-dependent effects of ZXH-2-107 and ZXH-8-004 on E abundance in a cell culture model of DENV infection. We infected Huh7.5 cells with DENV2 (MOI 1) and treated with increasing concentrations of ZXH-2-107, ZXH-2-107-Neg, ZXH-8-004, ZXH-8-004-Neg, GNF-2, and CVM-2-12-2 for 24 hours and then performed immunoblot analysis on cell lysates to monitor E protein abundance. As shown in Figures 5A and 5B, ZXH-8-004 was more effective than ZXH-2-107 at concentration-dependent depletion of E. Treatment of DENV2-infected Huh7.5 cells with ZXH-8-004 reduced intracellular E starting at a 1.25 μM dose. In contrast, ZXH-8-004-Neg and parental inhibitor CVM-2-12-2, which cannot induce degradation of E but can inhibit E during viral entry, did not exhibit an effect on E abundance until 10 μM (Figure 5C), consistent with the interpretation that this loss of E reflects general antiviral activity and not specific depletion of E. These data indicate that low concentrations of ZXH-8-004 (below 10 μM) yield CRBN-dependent E degradation, while high concentrations (10 μM or above) resulted in depletion of E that is CRBN-independent and may reflect inhibition of E other mechanisms or general antiviral activity.

**Figure 5.**
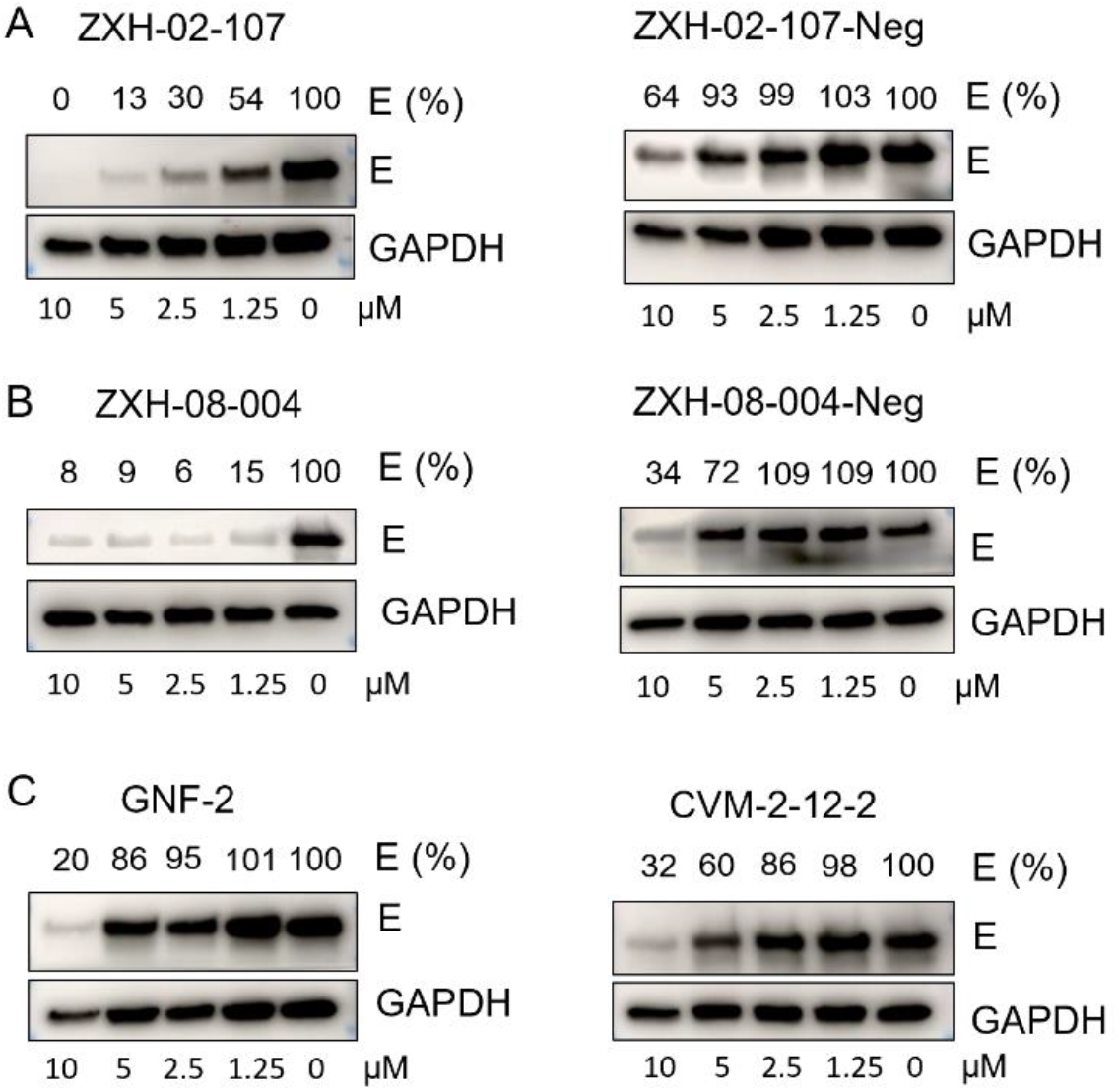
CRBN-dependence of ZXH-2-107- and ZXH-8-004-induced degradation of dengue E. A) Immunoblot analysis of E in DENV2-infected Huh 7.5 cells treated with ZXH-2-107 or ZXH-2-107-Neg for 24 h. B) Immunoblot analysis of E in DENV2-infected Huh 7.5 cells treated with ZXH-8-004 or ZXH-8-004-Neg for 24 h. C) Immunoblot analysis of E in DENV2-infected Huh 7.5 cells treated with GNF-2 or CVM-2-12-2 for 24 h. Semi-quantitative analysis was performed using the GAPDH signal for normalization.

To assess the general selectivity of degradation on a proteome-wide scale, we performed multiplexed mass spectrometry-based proteomics on MOLT4 cells treated with 3 μM of ZXH-2-107 and ZXH-8-004 for 5 hours, an experimental approach well-established for proteome-wide selectivity profiling of degraders.^37^ Both the mass spectrometry-based experiments and immunoblot analysis revealed no downregulation of ABL in cells treated with ZXH-8-004 and ZXH-2-107 (Figures 6A & S4). We did, however, observe downregulation of zinc-finger transcription factors such as ZFP91 and translation termination factor G1 to S phase transition 1 (GSPT1) for both compounds, all established off-targets of IMiD-based degraders.^38, 39^ ZXH-2-107 exhibited a higher number of off-targets compared to ZXH-8-004, potentially due to ZXH-2-107’s derivation from the kinase ligand GNF-2. Since GSPT1 is the primary off-target,^39^ and previous reports suggest that GSPT1 degradation can inhibit the replication of both RNA and DNA viruses,^40-42^ we sought to address the potential contribution of GSPT1 degradation to the antiviral activity of ZXH-8-004. We used a stable reporter cell line expressing a GSPT1-eGFP fusion protein fusion and mCherry reporter^43^ to detect depletion of GSPT1 by ZXH-8-004 after 24 hours, corresponding to the treatment duration used to evaluate ZXH-8-004 in DENV2-infected cells (Figure 3). ZXH-8-004 showed 80-fold less potent degradation of GSPT1 relative to CC-90009, a selective and potent GSPT1 degrader (Figure 6B).^44^

**Figure 6.**
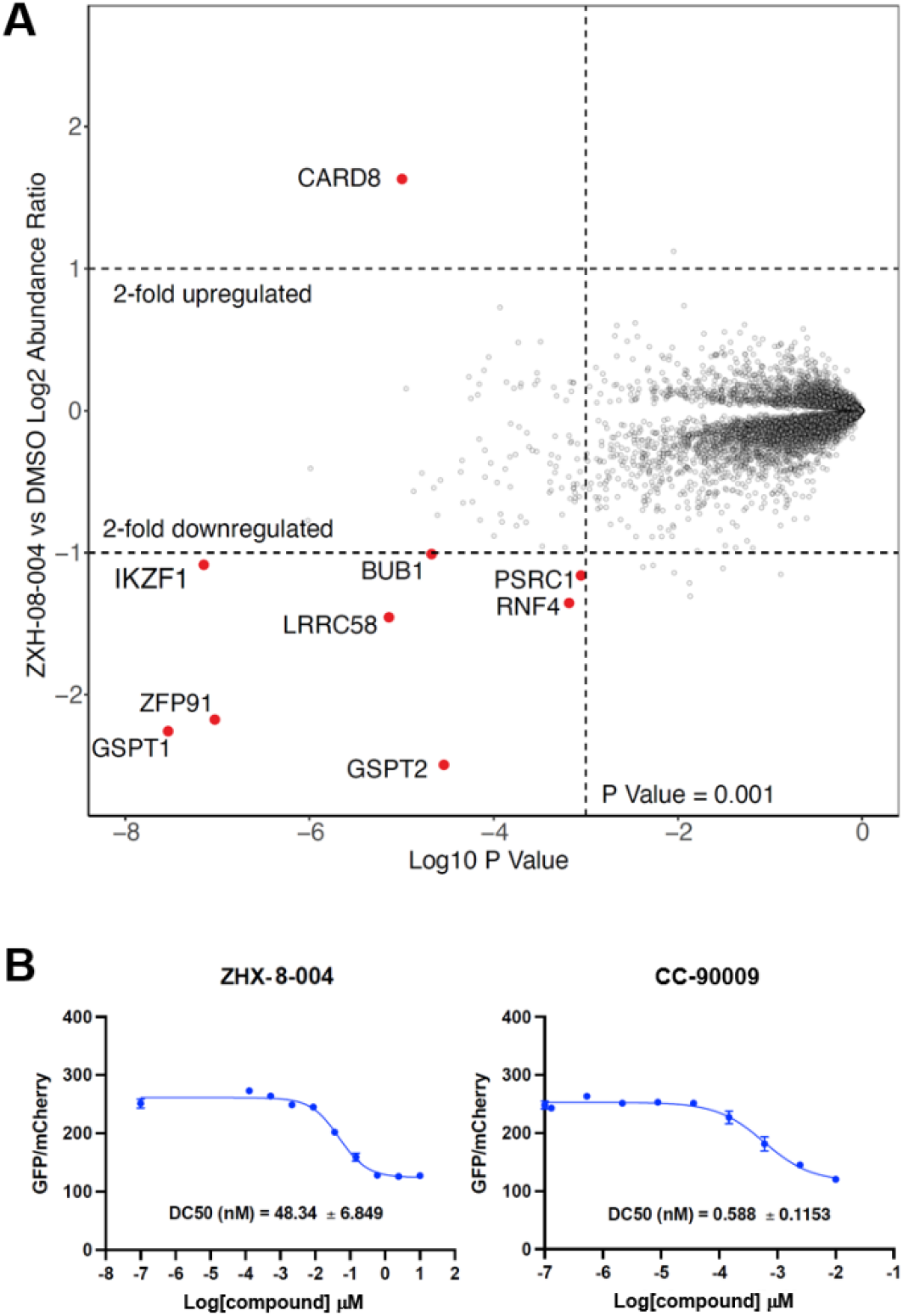
A) Proteome-wide degradation selectivity of ZXH-8-004 at a dose of 3 μM in MOLT4 cells after 5 hour treatment. B) Cellular GSPT1-induced degradation of ZXH-8-004 and CC-90009 assessed by reporter assay.

We then asked if GSPT1 degradation contributes to ZXH-8-004’s effect on E abundance and anti-DENV activity. We first investigated the effects of the selective GSPT1 degrader CC-90009 on the abundance of the E. This was achieved by treating Huh7.5 cells infected with DENV2 (MOI 1) with escalating concentrations of CC-90009 for 24 hours, followed by examining GSPT1 and E abundance by immunoblotting. As shown in Figure S5A, E levels decrease upon treatment with CC-90009, but this depletion is negligible compared to the extent of GSPT1 depletion. We next evaluated the antiviral activity of CC-90009 against infectious DENV2 in cell culture (Figure S5B). CC-90009’s potent degradation of GSPT1 was accompanied by weak antiviral activity, exhibiting ≤ 2-fold reduction in viral titer at concentrations that reduced GSPT1 to undetectable levels. These findings show that strong depletion of GSPT1 can have a very modest effect on E abundance but suggest that ZXH-8-004’s very modest effect on GSPT1 is unlikely to contribute to its effects on E abundance or to its antiviral activity.

### E degraders inhibit the four dengue serotypes

Having validated ZXH-2-107 and ZXH-8-004 as on-target E degraders, we next evaluated the compounds’ antiviral activity against infectious DENV2 in cell culture. Both ZXH-2-107 and ZXH-8-004 exhibit significantly enhanced antiviral activity compared to their parental inhibitors, GNF-2 and CVM-2-12-2, respectively (Figure 7 & S6). Since broad-spectrum activity across the DENV serotypes is of high interest, we additionally assessed the activity of the E PROTACs against representative strains of the other three DENV serotypes. Huh 7.5 cells were infected with each of the four DENV serotypes (DENV1-4) at a MOI=0.5, and the infected cells were then treated with E degraders at 5 and 2.5 μM concentrations. Both ZXH-2-107 and ZXH-8-004 demonstrate markedly increased antiviral efficacy across various serotypes compared to their respective parental inhibitors. This antiviral activity is CRBN-dependent, as ZXH-2-107-Neg and ZXH-8-004-Neg lack the boost in potency observed for the two degrader compounds (Figure 7 & S7).

**Figure 7.**
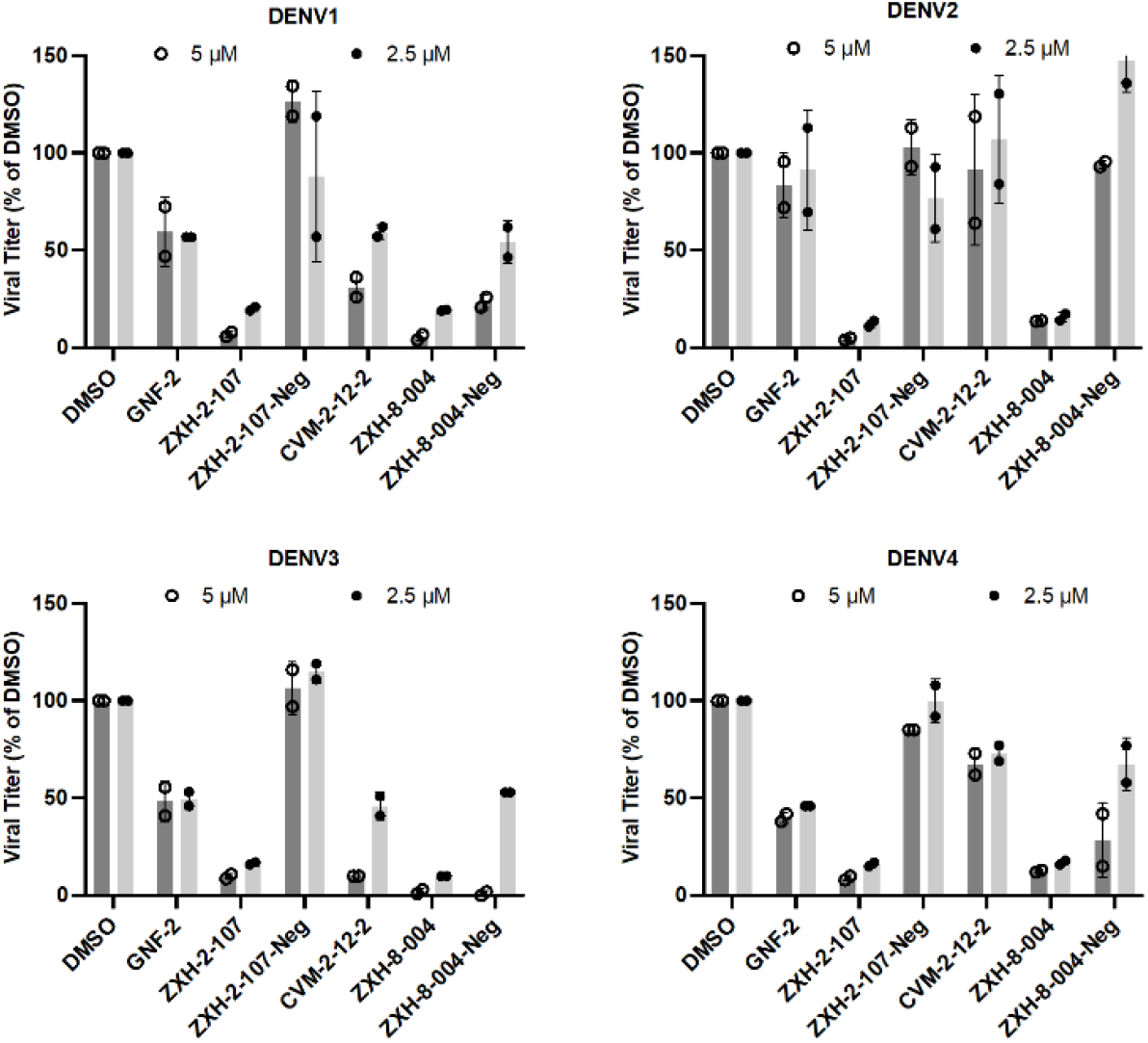
Antiviral activities of E degraders, negative controls, and parental inhibitors against DENV1, 2, 3, and 4 in Huh 7.5 cells at 2.5 μM and 5 μM. The yield of progeny virus in culture supernatants at 24 hours post-infection was quantified by viral plaque formation assay. Graphs show viral titer as % of the DMSO-treated, DENV2-infected controls.

### Conclusion

Traditional direct-acting small molecule antivirals target the functions of viral enzymes, such as polymerases and proteases. Small molecule antivirals against DENV that act against the RNA-dependent RNA polymerase activity of NS5^45^ or the nonstructural protein 4b (NS4B)^46^ have been successfully developed although none have yet been approved. Due to the occupancy-driven pharmacology of these compounds, they require very high affinity binding to their targets to prevent viral replication and the evolution of inhibitor-resistance. Since optimized degrader molecules can act catalytically, they can exert their effects at very low intracellular concentrations and may have superior resiliency to point mutations that reduce compound-binding, a phenomenon we observed for telaprevir-based PROTACs targeting the HCV NS3-4A protease.^22^ A major challenge, however, is that we currently do not know the repertoire of viral proteins that are susceptible to targeted protein degradation. The localization of many viral processes in specialized organelles may protect viral proteins from the cellular ubiquitin-proteasome machinery. In addition, many viruses have evolved mechanisms to ensure robust viral gene expression, raising questions about whether enough of the viral protein can be degraded to achieve significant antiviral activity. Here we explored whether previously reported small molecules that bind to the envelope (E) protein of DENV could be used as targeting moieties to generate bivalent degraders of this protein. Starting initially with GNF-2, we were able to generate ZXH-2-107, which causes CRL4^CRBN^-dependent degradation of E but retains inhibitory activity against ABL kinases. As inhibition of ABL kinase activity has been shown to inhibit DENV replication,^15^ we sought to generate an additional degrader that would only target E to provide a more precise pharmacological tool for assessing the antiviral effect of targeted degradation of E. To achieve this, we switched to an alternative E-ligand, CVM-2-12-2, that does not bind to ABL. Elaboration of this compound resulted in the development of ZXH-8-004, which effectively induces degradation of E without affecting ABL. A global proteomics experiment demonstrated that ZXH-8-004 has excellent selectivity for E with the only other significantly degraded proteins being the well-known ‘neosubstrates’ derived from the glutarimide recruiter (ZFP91 and GSPT1). We demonstrated that complete GSPT1 degradation induced by CC-90009 has very modest anti-DENV activity in our experimental model. Since ZHX-8-004 exhibits 80-fold less potent depletion of GSPT1 compared to CC-90009, this indictes that GSPT1 degradation by ZHX-8-004 may contribute marginally or not at all to its anti-DENV activity. We also showed that degradation requires engagement with the CRL4^CRBN^ E3 ligase in competition experiments in which pretreatment with lenalidomide rescued E abundance and in experiments using negative control compounds, ZXH-2-107-Neg and ZXH-8-004-Neg, in which the glutarimide amide carbonyl groups have been eliminated. These findings together demonstrate that dengue E is susceptible to targeted degradation mediated by the CRL4^CRBN^ ligase and that depletion of E from the cell by this pharmacological mechanism results in antiviral potency that exceeds that of the parental E inhibitors, which have occupancy-driven pharmacology.

Our finding that targeted degradation of dengue E is associated with significant antiviral activity raises several important questions regarding the mode(s) of action of this activity. E, like other viral glycoproteins, has important functions in both viral entry, at the beginning of the viral life cycle, and in viral particle production, near its end. Dengue virions first attach to the plasma membrane and then are internalized to the endosome until acidification of that compartment triggers E-mediated fusion of the viral and endosomal membranes, creating a pore through which the viral nucleocapsid can escape to the cytoplasm where the viral genome can be expressed. This process has been observed to occur on the order of minutes in live cell imaging studies,^47^ and it is unclear that the E3 CRL4^CRBN^ ubiquitin ligase could intercept and degrade E on the incoming virion prior to fusion. Post-fusion, E from the incoming virion is thought to remain associated with the endosome and is presumably degraded during endosome to lysosome maturation. Post-fusion E has no known further function for subsequent steps in the viral infectious cycle. Although targeted degradation of E could also affect the assembly of new virions, this mode of action has not been observed for classical inhibitors of dengue E or other viral envelope proteins.^48^ In addition, since dengue E has been thought to be co-translationally inserted through the ER membrane to the ER lumen, this raises questions as to how and where it is accessible and susceptible to the E3 CRL4^CRBN^ ubiquitin ligase and proteasome. Last but not least, since viral structural proteins are amongst the most abundantly expressed members of a viral proteome, they might be deemed unlikely targets for antiviral targeted protein degradation. Our results, however, indicate that this class of viral proteins are indeed susceptible to targeted protein degradation and that E degraders can exert potent antiviral activity against all four dengue serotypes. Future work to discover what steps (entry, particle assembly) in the viral life cycle are most affected by depletion of E and the sites of interaction of E with the host ubiquitin-proteasome system may provide valuable insights into the repertoire of viral proteins susceptible to targeted protein degradation. Relevant to our discovery, PROTAC-based degraders of the SARS-CoV-2 small envelope protein have also recently been proposed,^49^ but this idea has, to date, not been validated experimentally. Our findings provide important proof-of-concept that bivalent degraders can be generated to target viral envelope proteins, providing prototype drugs as starting points for the development and optimization of antiviral degraders as a new class of direct-acting antiviral drugs.

## Acknowledgements

We acknowledge funding from NIH/NIAID R01AI148632 and R01AI146152. Charles Rice (Rockefeller University) and Eva Harris (University of California, Berkeley) are gratefully acknowledged for sharing Huh7.5 and BHK21 cells, respectively. This work was supported by a Mark Foundation Emerging Leader Award 19-001-ELA (grant to E.S.F). Graphical abstract and schematic diagrams of the manuscript were created with BioRender.com.

