## Supporting Information-1 for "Discovery of Potent Degraders of the Dengue Virus Envelope Protein"

### Experimental Procedures

#### Chemistry

**General Methods.** Starting materials, reagents, and solvents were purchased from commercial suppliers and were used without further purification unless otherwise noted. All reactions were monitored using a Waters Acquity UPLC/MS system (Waters PDA eλ Detector, QDa Detector, Sample manager - FL, Binary Solvent Manager) using Acquity UPLC® BEH C18 column (2.1 x 50 mm, 1.7 µm particle size): solvent gradient = 85% A at 0 min, 1% A at 1.7 min; solvent A = 0.1% formic acid in Water; solvent B = 0.1% formic acid in Acetonitrile; flow rate : 0.6 mL/min. Reaction products were purified by flash column chromatography using CombiFlash®Rf with Teledyne Isco RediSep® normal-phase silica flash columns (4 g, 12 g, 24 g, 40 g or 80 g) and Waters HPLC system using SunFireTM Prep C18 column (19 x 100 mm, 5 µm particle size): solvent gradient = 80% A at 0 min, 10% A at 25 min; solvent A = 0.035% TFA in Water; solvent B = 0.035% TFA in MeOH; flow rate : 25 mL/min. $^{1}$H NMR spectra were recorded on 500 MHz Bruker Avance III spectrometers and $^{13}$C NMR spectra were recorded on 125 MHz Bruker Avance III spectrometer. Chemical shifts are reported in parts per million (ppm, δ) downfield from tetramethylsilane (TMS). Coupling constants (J) are reported in Hz. Spin multiplicities are described as br (broad), s (singlet), d (doublet), t (triplet), q (quartet), m (multiplet).

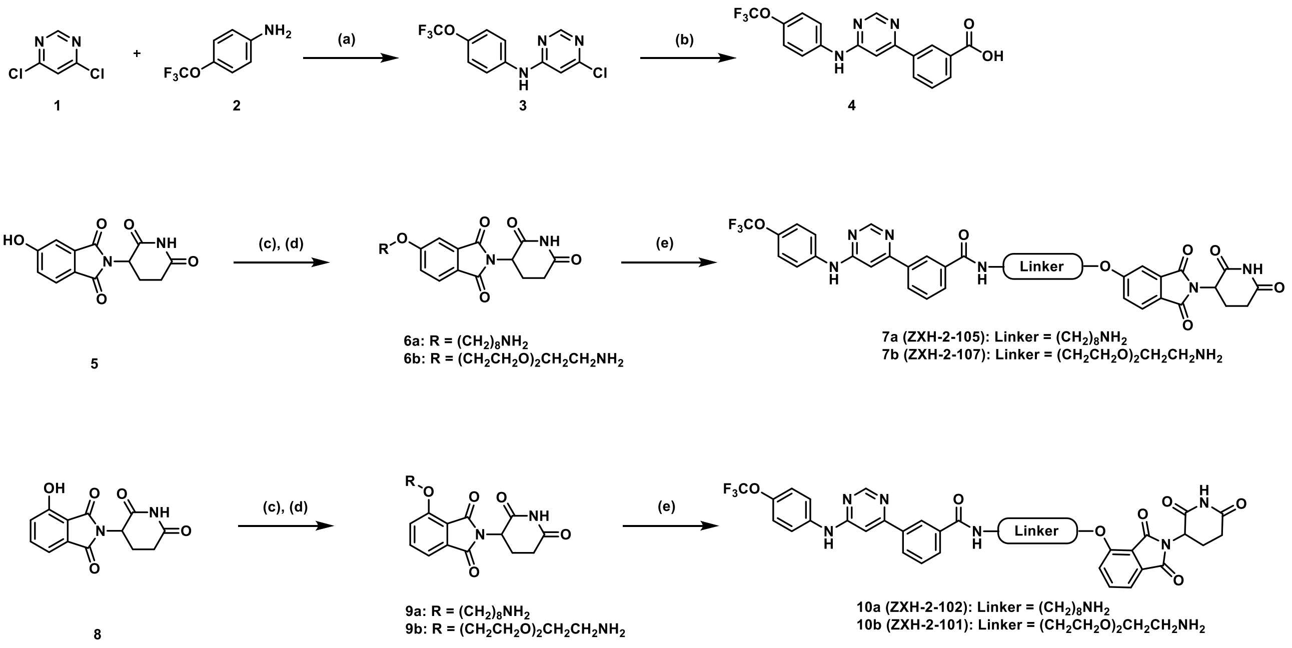


**Scheme S1.** Synthesis of GNF-2-CRBN compounds (a) DIEA, EtOH, 80 ºC (b) 3-boronobenzoic acid, Pd $(PhP_3)_4$, 2N $Na_2CO_3$, MeCN, 100 ºC (c) Br-R, $K_2CO_3$, DMF, 50 ºC (d) TFA, DCM, r.t. (e) **4**, HATU, DIEA, DMF, r.t.

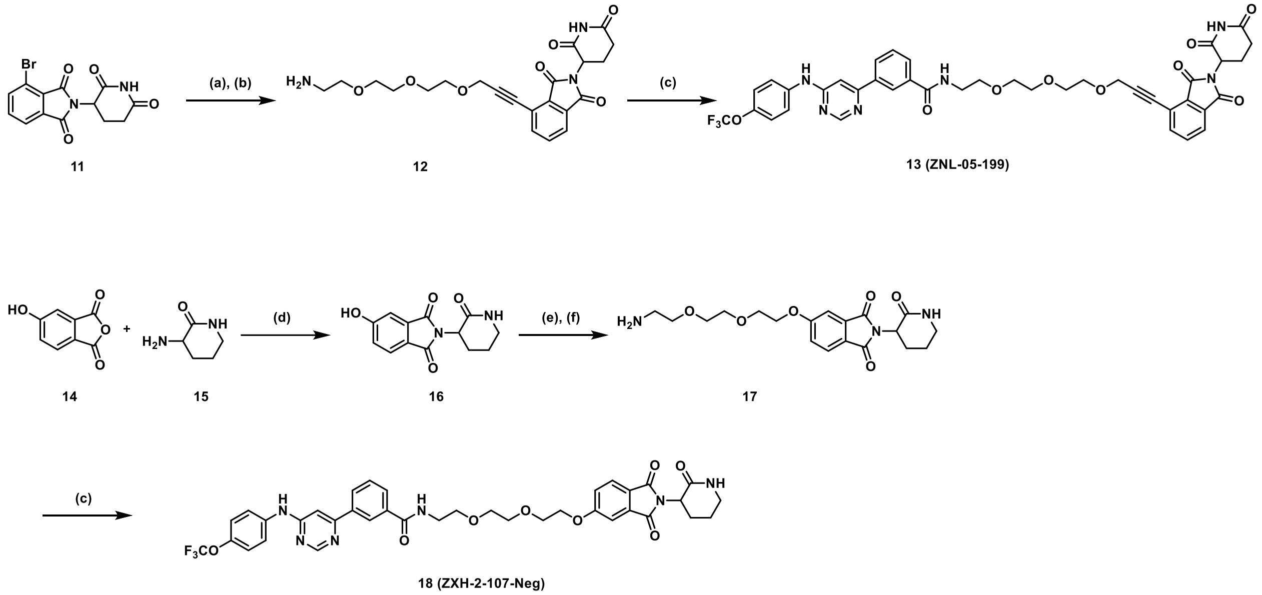


**Scheme S2.** Synthesis of GNF-2-CRBN compounds and negative control (a) Alkyne-(PEG)3-NHBoc, CuI, Pd $(PPh_3)_2Cl_2$, TEA, DMF, 80 ºC (b) TFA, DCM, r.t. (c) **4**, HATU, DIEA, DMF, r.t. (d) KOAc, AcOH, 90 ºC (e) Br-R, $K_2CO_3$, DMF, 50 ºC (f) TFA, DCM, r.t.

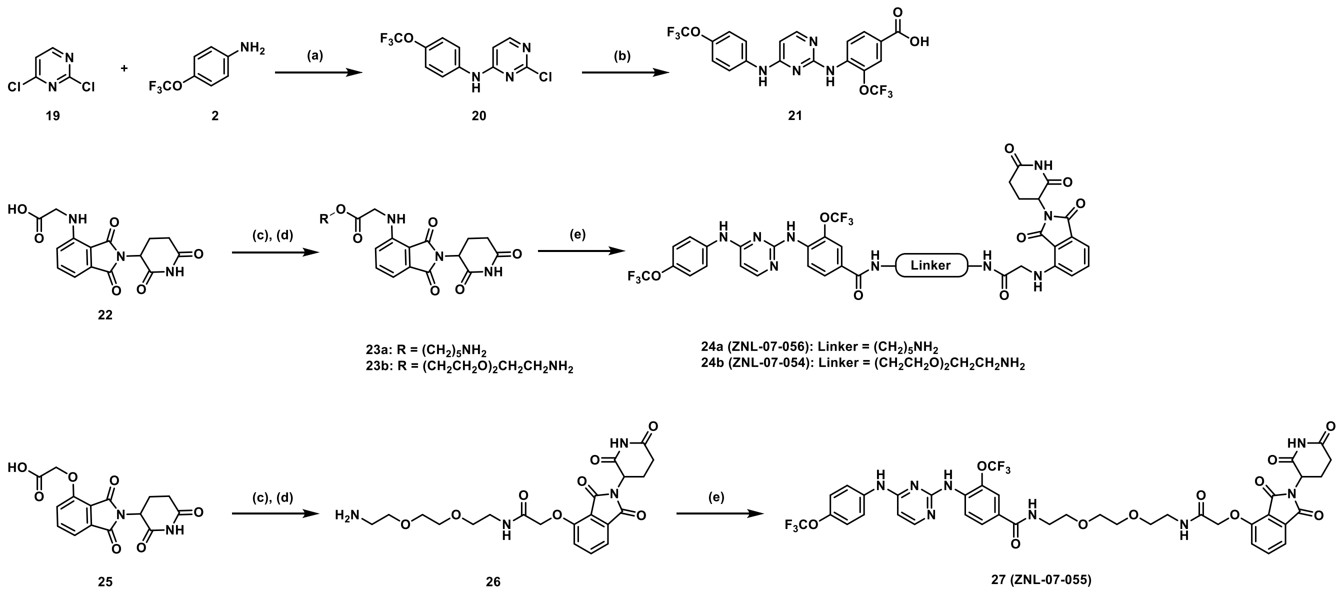


**Scheme S3.** Synthesis of CVM-2-12-2-CRBN compounds (a) DIEA, EtOH, 80 ºC (b) 4-amino-3-(trifluoromethoxy)benzoic acid, TFA, *s*-BuOH, 100 ºC (c) NHBoc-R, HATU, DIEA, DMF, r.t. (d) TFA, DCM, r.t. (e) **21**, HATU, DIEA, DMF, r.t.

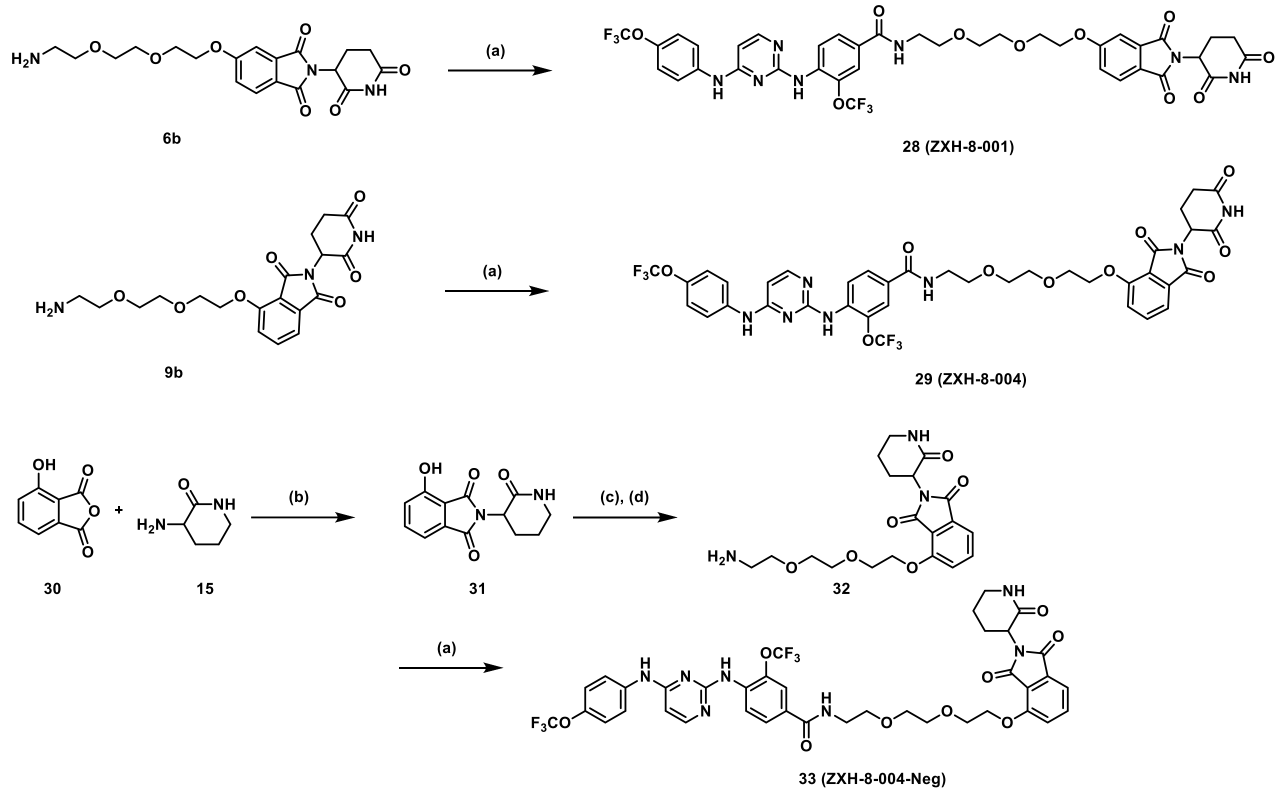


**Scheme S4.** Synthesis of CVM-2-12-2-CRBN compound and negative control (a) **21**, HATU, DIEA, DMF, r.t (b) KOAc, AcOH, 90 ºC (c) Br-PEG2-NHBoc, $K_2CO_3$, DMF, 50 ºC (d) TFA, DCM, r.t.

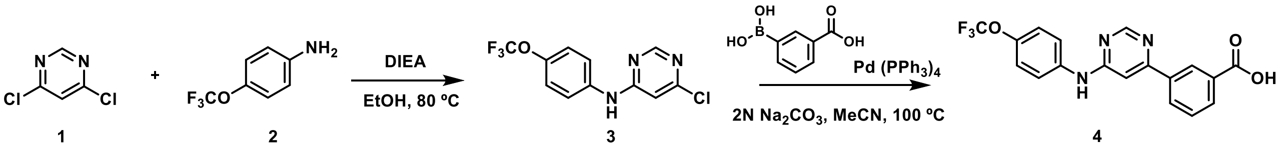


**Synthesis of 6-chloro-*N*-(4-(trifluoromethoxy)phenyl)pyrimidin-4-amine**

To the solution of 4,6-dichloro-pyrimidine (1.48 g, 10 mmol) and 4-trifluoromethoxyaniline (1.77 g, 10 mmol) in 50 mL of EtOH was added DIEA (1.9 mL, 11 mmol). The reaction mixture was stirred for 16 h at 80 ºC. Then the mixture was concentrated and purified by silica gel column chromatography (30% EA/Hexane) to give the title compound **3** (2.54 g, 88% yield). LC/MS (ESI) m/z calculated $[M+H]^+$ 290.02, found 290.23/292.23.

**Synthesis of 3-(6-((4-(trifluoromethoxy)phenyl)amino)pyrimidin-4-yl)benzoic acid**

To a mixture of 6-chloro-*N*-(4-(trifluoromethoxy)phenyl)pyrimidin-4-amine (1.5 g, 5.2 mmol), 3-carboxyphenylboronic acid (865 mg, 5.2 mmol), $Pd(PPh_3)_4$ (150 mg, 0.13 mmol), and 2N sodium carbonate (7.8 mL, 15.6 mmol) in acetonitrile/water (v/v = 1/1, 30 mL) was heated to 100 ºC under an nitrogen atmosphere. After refluxing for 6 h, the reaction mixture was filtered. The filtrate was cooled to room temperature and treated with 2N HCl (pH is around 4) to afford 3-(6-(4-(trifluoromethoxy) phenylamino)pyrimidin-4-yl)benzoic acid as the yellow precipitate, which was collected and washed with water and air-dried to give 1.46 g of the title compound (75% yield). LC/MS (ESI) m/z calculated $[M+H]^+$ 376.08, found 376.34.

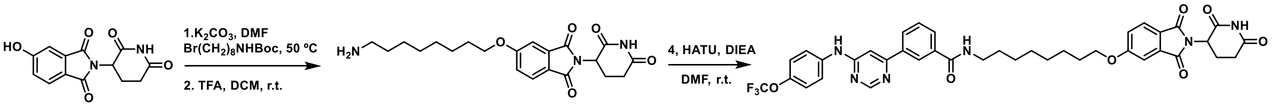


5

6a

ZXH-2-105

**Synthesis of 5-((8-aminooctyl)oxy)-2-(2,6-dioxopiperidin-3-yl)isoindoline-1,3-dione**

2-(2,6-dioxopiperidin-3-yl)-5-hydroxyisoindoline-1,3-dione (150mg, 0.55 mmol) was dissolved in DMF (3 mL), treated with $K_2CO_3$ (151 mg, 1.1 mmol), *tert*-butyl (8-bromooctyl)carbamate (169mg, 0.55 mmol) was added, and the mixture stirred 6 hours at 50 ºC. The reaction was diluted with water and extracted with EtOAc. Combined extracts were washed with brine, dried with $Na_2SO_4$, then concentrated and purified by silica gel chromatography to obtain *tert*-butyl (8-((2-(2,6-dioxopiperidin-3-yl)-1,3-dioxoisoindolin-5-yl)oxy)octyl)carbamate (222mg, 81%). LC/MS (ESI) m/z calculated $[M+H]^+$ 502.25, found [M+H-Boc] 402.28.

*tert*-butyl (8-((2-(2,6-dioxopiperidin-3-yl)-1,3-dioxoisoindolin-5-yl)oxy)octyl)carbamate (222mg, 0.44 mmol) was dissolved in 5 mL of DCM/TFA (1:1) and stirred for 2 hours at room temperature before being concentrated and dried to provide 5-((8-aminooctyl)oxy)-2-(2,6-dioxopiperidin-3-yl)isoindoline-1,3-dione. LC/MS (ESI) m/z calculated $[M+H]^+$ 402.20, found 402.32.

**Synthesis of *N*-(8-((2-(2,6-dioxopiperidin-3-yl)-1,3-dioxoisoindolin-5-yl)oxy)octyl)-3-(6-((4-(trifluoromethoxy)phenyl)amino)pyrimidin-4-yl)benzamide (ZXH-2-105)**

5-((8-aminooctyl)oxy)-2-(2,6-dioxopiperidin-3-yl)isoindoline-1,3-dione (90mg, 0.22 mmol) and 3-(6-(4- (trifluoromethoxy) phenylamino)pyrimidin-4-yl)benzoic acid (**4**) (83 mg, 0.22 mmol) were added to DMF (2.5 mL) followed by adding DIEA (193 µL, 1.1 mmol) and HATU (170 mg, 0.44 mmol), the reaction stirred for 30 minutes and then purified by HPLC to provide the title compound (56 mg, 0.073 mmol, 34%). $^1$H NMR (500 MHz, DMSO-$d_6$) δ 11.04 (s, 1H), 10.06 (s, 1H), 8.73 (d, J = 1.0 Hz, 1H), 8.56 (t, J = 5.6 Hz, 1H), 8.42 (t, J = 1.9 Hz, 1H), 8.09 (dt, J = 8.0, 1.4 Hz, 1H), 7.93 (dt, J = 7.8, 1.5 Hz, 1H), 7.80 – 7.76 (m, 2H), 7.74 (d, J = 8.3 Hz, 1H), 7.57 (t, J = 7.8 Hz, 1H), 7.33 (d, J = 2.3 Hz, 1H), 7.31 (d, J = 8.6 Hz, 2H), 7.28 – 7.23 (m, 2H), 5.04 (dd, J = 12.8, 5.5 Hz, 1H), 4.09 (t, J = 6.5 Hz, 2H), 3.23 (q, J = 6.7 Hz, 2H), 2.82 (ddd, J = 16.9, 13.8, 5.4 Hz, 1H), 2.52 (dt, J = 17.1, 3.0 Hz, 1H), 2.01 – 1.95 (m, 1H), 1.73 – 1.64 (m, 2H), 1.48 (q, J = 6.8 Hz, 2H), 1.35 (q, J = 6.9 Hz, 2H), 1.30 – 1.23 (m, 6H). $^{13}$C NMR (125 MHz, DMSO) δ 173.24, 170.41, 167.36, 167.28, 166.09, 164.57, 161.41, 160.23, 159.12, 158.54, 158.13, 143.69, 139.21, 136.48, 135.93, 134.41, 129.66, 129.52, 129.46, 126.01, 125.75, 123.32, 122.23, 121.72, 121.68,

121.17, 119.64, 109.27, 103.27, 69.27, 49.43, 31.42, 29.52, 29.17, 29.08, 28.83, 26.89, 25.79, 22.54.LC/MS (ESI) m/z calculated $[M+H]^+$ 759.27, found 759.20.

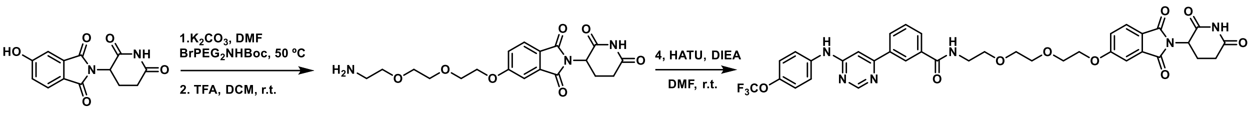


5 6b ZXH-2-107

**Synthesis of *N*-(2-(2-(2-((2-(2,6-dioxopiperidin-3-yl)-1,3-dioxoisoindolin-5-yl)oxy)ethoxy)ethoxy)ethyl)-3-(6-((4-(trifluoromethoxy)phenyl)amino)pyrimidin-4-yl)benzamide (ZXH-2-107)**

By using 5-(2-(2-(2-aminoethoxy)ethoxy)ethoxy)-2-(2,6-dioxopiperidin-3-yl)isoindoline-1,3-dione (**6b**) and 3-(6-(4- (trifluoromethoxy) phenylamino)pyrimidin-4-yl)benzoic acid (**4**) following the same procedure as **ZXH-2-105** was prepared as a yellow solid (35 mg, 0.045 mmol, 18% two steps). $^1$H NMR (500 MHz, DMSO-$d_6$) δ 11.11 (s, 1H), 10.12 (s, 1H), 8.80 (d, J = 1.0 Hz, 1H), 8.71 (t, J = 5.6 Hz, 1H), 8.51 (t, J = 1.8 Hz, 1H), 8.16 (dt, J = 7.8, 1.4 Hz, 1H), 8.01 (dt, J = 7.8, 1.4 Hz, 1H), 7.87 – 7.83 (m, 2H), 7.80 (d, J = 8.3 Hz, 1H), 7.64 (t, J = 7.8 Hz, 1H), 7.41 (d, J = 2.3 Hz, 1H), 7.40 – 7.36 (m, 2H), 7.34 – 7.31 (m, 2H), 5.11 (dd, J = 12.8, 5.4 Hz, 1H), 4.29 – 4.25 (m, 2H), 3.82 – 3.76 (m, 2H), 3.66 – 3.56 (m, 6H), 3.47 (q, J = 5.8 Hz, 2H), 2.89 (ddd, J = 16.8, 13.7, 5.4 Hz, 1H), 2.63 – 2.52 (m, 2H), 2.08 – 2.01 (m, 1H). $^{13}$C NMR (125 MHz, DMSO) δ 173.24, 170.41, 167.31, 167.25, 166.32, 164.35, 161.40, 160.20, 158.53, 158.13, 143.70, 139.20, 136.52, 135.61, 134.36, 129.67, 129.59, 129.55, 126.06, 125.71, 123.52, 122.24, 121.73, 121.31, 119.65, 117.25, 114.93, 109.29, 103.26, 70.35, 70.09, 69.33, 69.10, 68.87, 49.43, 31.41, 22.53. LC/MS (ESI) m/z calculated $[M+H]^+$ 763.23, found 763.21.

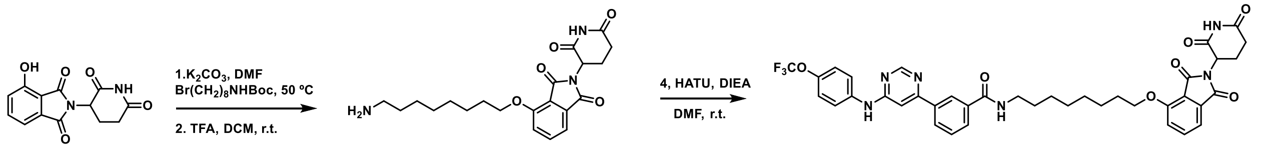


8 9a ZXH-2-102

**Synthesis of *N*-(8-((2-(2,6-dioxopiperidin-3-yl)-1,3-dioxoisoindolin-4-yl)oxy)octyl)-3-(6-((4-(trifluoromethoxy)phenyl)amino)pyrimidin-4-yl)benzamide (ZXH-2-102)**

Compound **ZXH-2-102** was synthesized by starting with 2-(2,6-dioxopiperidin-3-yl)-4-hydroxyisoindoline-1,3-dione (**8**) and linker *tert*-butyl (8-bromooctyl)carbamate following the same procedure as **ZXH-2-105** was prepared as a yellow solid (12 mg, 0.016 mmol, 23 % two steps). $^1$H NMR (500 MHz, DMSO-$d_6$) δ 11.10 (s, 1H), 9.94 (s, 1H), 8.77 (d, J = 1.1 Hz, 1H), 8.62 (t, J = 5.6 Hz, 1H), 8.50 (t, J = 1.8 Hz, 1H), 8.17 (dt, J = 7.8, 1.3 Hz, 1H), 7.97 (dt, J = 7.8, 1.3 Hz, 1H), 7.88 – 7.83 (m, 2H), 7.80 (dd, J = 8.5, 7.3 Hz, 1H), 7.63 (t, J = 7.8 Hz, 1H), 7.50 (d, J = 8.5 Hz, 1H), 7.43 (d, J = 7.2 Hz, 1H), 7.36 (d, J = 8.2 Hz, 1H), 7.33 (d, J = 1.2 Hz, 1H), 5.08 (dd, J = 12.8, 5.4 Hz, 1H), 4.19 (t, J = 6.4 Hz, 2H), 3.68 – 3.57 (m, 2H), 3.29 (q, J = 6.7 Hz, 2H), 2.88 (ddd, J = 17.0, 13.9, 5.5 Hz, 1H), 2.63 – 2.52 (m, 2H), 2.05 – 2.00 (m, 1H), 1.79 – 1.73 (m, 2H), 1.56 (t, J = 6.7 Hz, 2H), 1.47 (t, J = 7.5 Hz, 2H), 1.38 – 1.33 (m, 6H). $^{13}$C NMR (125 MHz, DMSO) δ 173.28, 170.45, 167.34, 166.24, 165.79, 161.29, 161.27, 158.76, 156.49, 143.39, 139.57, 137.51, 137.32, 135.88, 133.70, 129.45, 129.35, 125.84, 122.20, 121.69, 121.36, 120.24, 119.66, 116.66,

115.60, 103.08, 69.27, 49.20, 31.41, 29.51, 29.18, 29.11, 28.88, 26.90, 25.73, 22.47. LC/MS (ESI) m/z calculated $[M+H]^+$ 759.27, found 759.30.

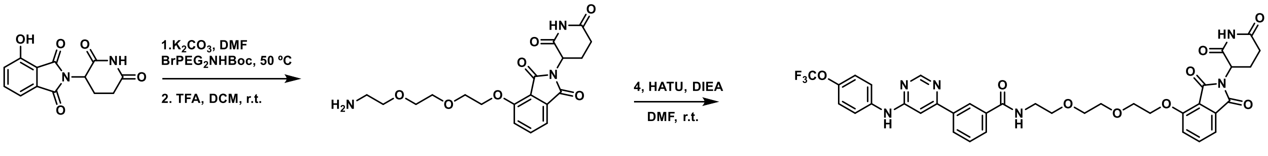


8 9b ZXH-2-101

**Synthesis of *N*-(2-(2-(2-((2-(2,6-dioxopiperidin-3-yl)-1,3-dioxoisoindolin-4-yl)oxy)ethoxy)ethoxy)ethyl)-3-(6-((4-(trifluoromethoxy)phenyl)amino)pyrimidin-4-yl)benzamide (ZXH-2-101)**

Compound **ZXH-2-101** was synthesized by starting with 2-(2,6-dioxopiperidin-3-yl)-4-hydroxyisoindoline-1,3-dione (**8**) and linker *tert*-butyl (2-(2-(2-bromoethoxy)ethoxy)ethyl)carbamate following the same procedure as **ZXH-2-105** was prepared as a yellow solid (14 mg, 0.018 mmol, 26 % two steps). $^1$H NMR (500 MHz, DMSO-$d_6$) δ 11.03 (s, 1H), 10.03 (s, 1H), 8.72 (d, J = 1.0 Hz, 1H), 8.63 (t, J = 5.6 Hz, 1H), 8.43 (t, J = 1.8 Hz, 1H), 8.09 (dt, J = 7.6, 1.4 Hz, 1H), 7.93 (dt, J = 7.8, 1.4 Hz, 1H), 7.81 – 7.74 (m, 2H), 7.71 (dd, J = 8.5, 7.3 Hz, 1H), 7.56 (t, J = 7.7 Hz, 1H), 7.41 (d, J = 8.5 Hz, 1H), 7.36 (d, J = 7.2 Hz, 1H), 7.30 (d, J = 8.1 Hz, 1H), 7.26 (s, 1H), 5.01 (dd, J = 12.8, 5.5 Hz, 1H), 4.27 – 4.21 (m, 2H), 3.76 – 3.71 (m, 2H), 3.60 (dd, J = 5.9, 3.6 Hz, 2H), 3.54 – 3.48 (m, 4H), 3.39 (q, J = 5.8 Hz, 2H), 2.80 (ddd, J = 16.9, 13.8, 5.4 Hz, 1H), 2.59 – 2.44 (m, 2H), 2.00 – 1.92 (m, 1H). $^{13}$C NMR (125 MHz, DMSO) δ 173.25, 170.41, 167.26, 166.33, 165.75, 161.40, 160.30, 158.80, 158.52, 158.23, 158.20, 156.27, 143.65, 139.25, 137.42, 136.60, 135.61, 133.70, 129.65, 129.57, 129.54, 126.04, 122.23, 121.68, 120.43, 119.65, 116.78, 115.85, 103.26, 70.57, 70.17, 69.33, 69.31, 69.17, 49.22, 31.42, 22.46.LC/MS (ESI) m/z calculated $[M+H]^+$ 763.23, found 763.12.

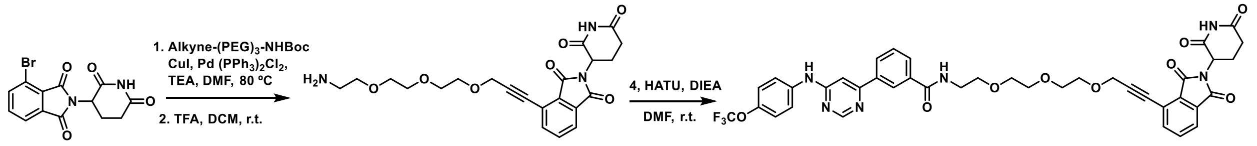


11 12 13 (ZNL-05-199)

**Synthesis of 4-(3-(2-(2-(2-aminoethoxy)ethoxy)ethoxy)prop-1-yn-1-yl)-2-(2,6-dioxopiperidin-3-yl)isoindoline-1,3-dione (12)**

A mixture of 4-bromo-2-(2,6-dioxopiperidin-3-yl)isoindoline-1,3-dione (**11**) (100 mg, 0.3 mmol), *tert*-butyl (2-(2-(2-(prop-2-yn-1-yloxy)ethoxy)ethoxy)ethyl)carbamate (86 mg, 0.3 mmol), $Pd(PPh_3)_2Cl_2$ (21 mg, 0.03 mmol), CuI (11 mg, 0.06 mmol), and TEA(0.5 mL) in DMF(2 mL) then stirred at 80 °C for 6 h under the nitrogen atmosphere. The reaction mixture was diluted with water (10 mL) and extracted with ethyl acetate (15 mL × 3). The organic layers were combined and washed with brine, dried over $Na_2SO_4$, filtered, and concentrated. The residue obtained was purified with column chromatography (10% MeOH/DCM) to yield intermediate *tert*-butyl (2-(2-(2-((3-(2-(2,6-dioxopiperidin-3-yl)-1,3-dioxoisoindolin-4-yl)prop-2-yn-1-

yl)oxy)ethoxy)ethoxy)ethyl)carbamate as a yellow solid (52 mg, 32%). LC/MS (ESI) m/z calculated $[M+H]^+$ 544.22, found 544.23.

*tert*-butyl (2-(2-(2-((3-(2-(2,6-dioxopiperidin-3-yl)-1,3-dioxoisoindolin-4-yl)prop-2-yn-1-yl)oxy)ethoxy)ethoxy)ethyl)carbamate carbamate (52mg, 0.096 mmol) was dissolved in 4 mL of DCM/TFA (1:1) and stirred for 2 hours at room temperature before being concentrated and dried to provide 4-(3-(2-(2-(2-aminoethoxy)ethoxy)ethoxy)prop-1-yn-1-yl)-2-(2,6-dioxopiperidin-3-yl)isoindoline-1,3-dione (**12**). LC/MS (ESI) m/z calculated $[M+H]^+$ 444.17, found 444.20.

**Synthesis of *N*-(2-(2-(2-((3-(2-(2,6-dioxopiperidin-3-yl)-1,3-dioxoisoindolin-4-yl)prop-2-yn-1-yl)oxy)ethoxy)ethoxy)ethyl)-3-(6-((4-(trifluoromethoxy)phenyl)amino)pyrimidin-4-yl)benzamide (ZNL-05-199)**

4-(3-(2-(2-(2-aminoethoxy)ethoxy)ethoxy)prop-1-yn-1-yl)-2-(2,6-dioxopiperidin-3-yl)isoindoline-1,3-dione (40mg, 0.09 mmol) and 3-(6-(4- (trifluoromethoxy) phenylamino)pyrimidin-4-yl)benzoic acid (**4**) (34 mg, 0.09 mmol) were added to DMF (2.0 mL) followed by adding DIEA (72 µL, 0.45 mmol) and HATU (68 mg, 0.18 mmol), the reaction stirred for 30 minutes and then purified by HPLC to provide the title compound (26 mg, 0.033 mmol, 37%). $^1$H NMR (500 MHz, DMSO-$d_6$) δ 11.14 (s, 1H), 8.80 (s, 1H), 8.71 (t, J = 5.6 Hz, 1H), 8.51 (t, J = 1.9 Hz, 1H), 8.17 (dt, J = 8.0, 1.4 Hz, 1H), 8.01 (dt, J = 7.8, 1.4 Hz, 1H), 7.93 (s, 1H), 7.91 (s, 2H), 7.89 – 7.81 (m, 2H), 7.37 (d, J = 8.6 Hz, 2H), 7.33 (s, 1H), 5.17 (dd, J = 12.9, 5.5 Hz, 1H), 4.45 (s, 2H), 3.65 (dd, J = 5.8, 3.1 Hz, 2H), 3.62 – 3.54 (m, 8H), 3.47 (q, J = 5.9 Hz, 2H), 2.89 (ddd, J = 17.1, 13.8, 5.4 Hz, 1H), 2.65 – 2.51 (m, 2H), 2.49 (d, J = 6.4 Hz, 2H), 2.06 (ddq, J = 11.1, 5.5, 2.8 Hz, 1H). $^{13}$C NMR (125 MHz, DMSO) δ 173.25, 170.23, 166.92, 166.82, 166.41, 161.38, 160.35, 158.79, 158.51, 158.20, 143.67, 139.20, 138.08, 136.62, 135.58, 132.22, 131.00, 129.64, 129.61, 129.56, 128.68, 126.23, 126.05, 124.25, 122.23, 121.70, 119.63, 103.23, 91.22, 84.57, 70.19, 70.09, 70.05, 69.34, 69.29, 58.51, 49.58, 31.35, 22.37. LC/MS (ESI) m/z calculated $[M+H]^+$ 801.24, found 801.39.

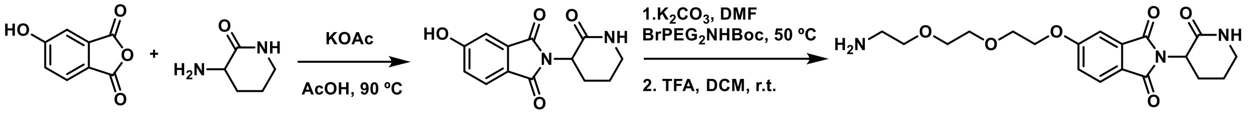


**Synthesis of 5-(2-(2-(2-aminoethoxy)ethoxy)ethoxy)-2-(2-oxopiperidin-3-yl)isoindoline-1,3-dione (17)**

A mixture of 5-hydroxyisobenzofuran-1,3-dione (1g , 6.1 mmol), 3-aminopiperidin-2-one hydrochloride (1g , 6.7 mmol), and NaOAc (1.8g , 18.6 mmol) was dissolved in HOAc (20 mL), and the resulting mixture was stirred at 90 °C for 16 h. After the reaction was completed, the solution was filtered and the filtration residue was washed by water and dried to yield a yellow solid (**16**), which was used in the next step without further purification (870 mg, 55%).

5-hydroxy-2-(2-oxopiperidin-3-yl)isoindoline-1,3-dione (50mg, 0.17 mmol) was dissolved in DMF (1.5 mL), treated with $K_2CO_3$ (46 mg, 0.34 mmol), *tert*-butyl (2-(2-(2-bromoethoxy)ethoxy)ethyl)carbamate (55 mg, 0.17 mmol) was added, and the mixture stirred 6 hours at 50 ºC. The reaction was diluted with water and extracted with EtOAc. Combined extracts were washed with brine, dried with $Na_2SO_4$, then concentrated and purified by silica gel chromatography to obtain *tert*-butyl (2-(2-(2-((1,3-dioxo-2-(2-oxopiperidin-3-yl)isoindolin-5-

yl)oxy)ethoxy)ethoxy)ethyl)carbamate (31 mg, 38%). LC/MS (ESI) m/z calculated $[M+H]^+$ 492.23, found [M+H-Boc] 392.45

*tert*-butyl (2-(2-(2-((1,3-dioxo-2-(2-oxopiperidin-3-yl)isoindolin-5-yl)oxy)ethoxy)ethoxy)ethyl)carbamate (31mg, 0.063 mmol) was dissolved in 2 mL of DCM/TFA (1:1) and stirred for 2 hours at room temperature before being concentrated and dried to provide 5-(2-(2-(2-aminoethoxy)ethoxy)ethoxy)-2-(2-oxopiperidin-3-yl)isoindoline-1,3-dione (**17**). LC/MS (ESI) m/z calculated $[M+H]^+$ 392.17, found 392.45

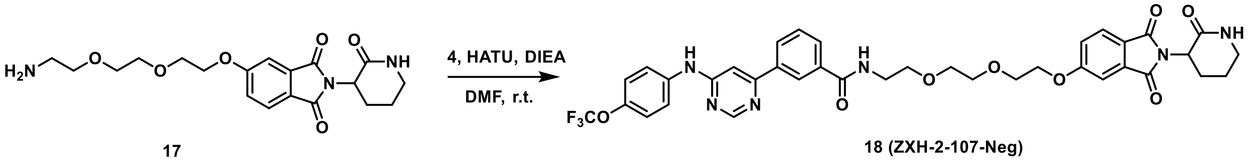


**Synthesis of *N*-(2-(2-(2-((1,3-dioxo-2-(2-oxopiperidin-3-yl)isoindolin-5-yl)oxy)ethoxy)ethoxy)ethyl)-3-(6-((4-(trifluoromethoxy)phenyl)amino)pyrimidin-4-yl)benzamide (ZXH-2-107-Neg)**

5-(2-(2-(2-aminoethoxy)ethoxy)ethoxy)-2-(2-oxopiperidin-3-yl)isoindoline-1,3-dione (30mg, 0.076 mmol) and 3-(6-(4- (trifluoromethoxy) phenylamino)pyrimidin-4-yl)benzoic acid (**4**) (29 mg, 0.064 mmol) were added to DMF (2.0 mL) followed by adding DIEA (66 µL, 0.38 mmol) and HATU (60 mg, 0.152 mmol), the reaction stirred for 30 minutes and then purified by HPLC to provide the title compound (26 mg, 32%). $^1$H NMR (500 MHz, DMSO-$d_6$) δ 9.92 (s, 1H), 8.76 (d, J = 1.1 Hz, 1H), 8.70 (t, J = 5.6 Hz, 1H), 8.52 (t, J = 1.8 Hz, 1H), 8.18 (dt, J = 7.8, 1.5 Hz, 1H), 7.99 (dt, J = 7.8, 1.4 Hz, 1H), 7.86 (d, J = 2.2 Hz, 1H), 7.84 (d, J = 3.2 Hz, 2H), 7.76 (s, 1H), 7.63 (t, J = 7.8 Hz, 1H), 7.37 (d, J = 5.5 Hz, 1H), 7.36 (d, J = 1.7 Hz, 2H), 7.32 (d, J = 1.2 Hz, 1H), 7.29 (dd, J = 8.3, 2.4 Hz, 1H), 4.56 (dd, J = 12.0, 6.3 Hz, 1H), 4.27 – 4.24 (m, 2H), 3.81 – 3.76 (m, 2H), 3.66 – 3.56 (m, 6H), 3.47 (q, J = 5.9 Hz, 2H), 3.28 – 3.15 (m, 2H), 2.21 (qd, J = 12.2, 4.8 Hz, 1H), 2.02 – 1.94 (m, 1H), 1.93 – 1.78 (m, 2H). $^{13}$C NMR (125 MHz, DMSO) δ 167.53, 167.50, 166.41, 164.17, 161.28, 161.17, 158.74, 143.37, 143.35, 139.58, 137.32, 135.57, 134.50, 129.46, 125.89, 125.45, 123.71, 122.20, 121.69, 121.34, 121.03, 119.66, 109.09, 103.07, 70.34, 70.09, 69.32, 69.11, 68.81, 49.48, 41.84, 26.22, 22.14.LC/MS (ESI) m/z calculated $[M+H]^+$ 749.23, found 749.78.

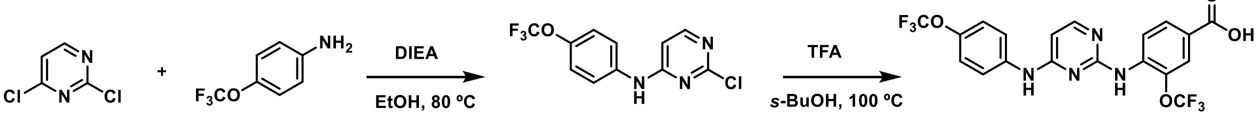


**Synthesis of 2-chloro-*N*-(4-(trifluoromethoxy)phenyl)pyrimidin-4-amine**

To the solution of 2,4-dichloropyrimidine (1.48 g, 10 mmol) and 4-trifluoromethoxyaniline (1.77 g, 10 mmol) in 50 mL of EtOH was added DIEA (1.9 mL, 11 mmol). The reaction mixture was stirred for 16 h at 80 ºC. Then the mixture was concentrated and purified by silica gel column chromatography (30% EA/Hexane) to give the title compound **20** (2.4 g, 83% yield). LC/MS (ESI) m/z calculated $[M+H]^+$ 290.02, found 290.24/292.19

**Synthesis of 4-((4-((4-(trifluoromethoxy)phenyl)amino)pyrimidin-2-yl)amino)-3-(trifluoromethyl)benzoic acid**

To a mixture of 2-chloro-*N*-(4-(trifluoromethoxy)phenyl)pyrimidin-4-amine (550 mg, 19 mmol), 4-amino-3-(trifluoromethoxy)benzoic acid (420 mg, 1.9 mmol), and TFA (190 µL, 3.8 mmol) in *s*-butanol (6 mL) was heated to 100 ºC. After refluxing for 6 h, the reaction was diluted with water and extracted with EtOAc. Combined extracts were washed with brine, dried with $Na_2SO_4$, then concentrated and purified by silica gel chromatography to obtain 4-((4-((4-(trifluoromethoxy)phenyl)amino)pyrimidin-2-yl)amino)-3-(trifluoromethyl)benzoic acid (380 mg, 42%).LC/MS (ESI) m/z calculated $[M+H]^+$ 475.08, found 475.42.

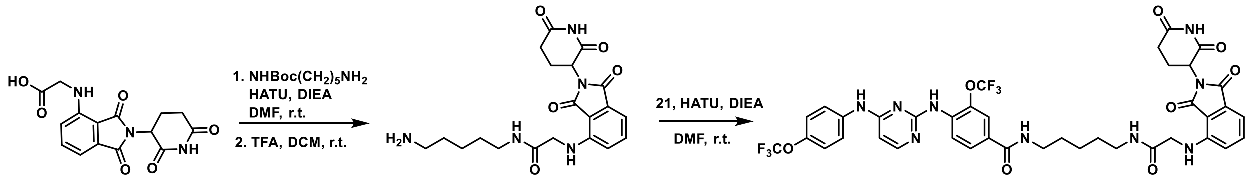


22 23a ZNL-07-056

**Synthesis of *N*-(5-aminopentyl)-2-((2-(2,6-dioxopiperidin-3-yl)-1,3-dioxoisoindolin-4-yl)amino)acetamide (23a)**

(2-(2,6-dioxopiperidin-3-yl)-1,3-dioxoisoindolin-4-yl)glycine (17mg, 0.05 mmol) and *tert*-butyl (5-aminopentyl)carbamate (10 mg, 0.05 mmol) were added to DMF (1.5 mL) followed by adding DIEA (43 µL, 0.25 mmol) and HATU (38 mg, 0.1 mmol), the reaction stirred for 30 minutes and then purified by silica gel chromatography(10% MeOH in DCM) to provide the title compound (25 mg, 98%). LC/MS (ESI) m/z calculated $[M+H]^+$ 516.24, found [M+H-Boc] 416.45.

*tert*-butyl (5-(2-((2-(2,6-dioxopiperidin-3-yl)-1,3-dioxoisoindolin-4-yl)amino)acetamido)pentyl)carbamate (23mg, 0.044 mmol) was dissolved in 2 mL of DCM/TFA (1:1) and stirred for 2 hours at room temperature before being concentrated and dried to provide *N*-(5-aminopentyl)-2-((2-(2,6-dioxopiperidin-3-yl)-1,3-dioxoisoindolin-4-yl)amino)acetamide(**23a**). LC/MS (ESI) m/z calculated $[M+H]^+$ 416.19, found 416.45.

**Synthesis of *N*-(5-(2-((2-(2,6-dioxopiperidin-3-yl)-1,3-dioxoisoindolin-4-yl)amino)acetamido)pentyl)-3-(trifluoromethoxy)-4-((4-((4-(trifluoromethoxy)phenyl)amino)pyrimidin-2-yl)amino)benzamide (ZNL-07-056)**

*N*-(5-aminopentyl)-2-((2-(2,6-dioxopiperidin-3-yl)-1,3-dioxoisoindolin-4-yl)amino)acetamide (20mg, 0.048 mmol) and 4-((4-((4-(trifluoromethoxy)phenyl)amino)pyrimidin-2-yl)amino)-3-(trifluoromethyl)benzoic acid (**21**) (23 mg, 0.048 mmol) were added to DMF (1.0 mL) followed by adding DIEA (42 µL, 0.24 mmol) and HATU (36 mg, 0.096 mmol), the reaction stirred for 30 minutes and then purified by HPLC to provide the title compound (16 mg, 38%).$^{1}$H NMR (500 MHz, DMSO-$d_6$) δ 11.11 (s, 1H), 9.64 (s, 1H), 8.81 (s, 1H), 8.53 (t, J = 5.6 Hz, 1H), 8.14 – 8.08 (m, 2H), 8.06 (d, J = 5.7 Hz, 1H), 7.85 (d, J = 7.6 Hz, 2H), 7.77 – 7.72 (m, 2H), 7.59 (dd, J = 8.5, 7.1 Hz, 1H), 7.24 (d, J = 8.6 Hz, 2H), 7.07 (d, J = 7.1 Hz, 1H), 6.95 (t, J = 5.6 Hz, 1H), 6.86 (d, J = 8.6 Hz, 1H), 6.32 (d, J = 5.8 Hz, 1H), 5.08 (dd, J = 12.8, 5.4 Hz, 1H), 3.92 (d, J = 5.7 Hz, 2H), 3.26 (q, J = 6.7 Hz, 2H), 3.12 (q, J = 6.6 Hz, 2H), 2.90 (ddd, J = 16.8, 13.7, 5.2 Hz, 1H), 2.63 – 2.52 (m, 2H), 2.10 – 1.99 (m, 1H), 1.50 (dp, J = 37.9, 7.3 Hz, 4H), 1.36 – 1.26 (m, 2H). $^{13}$C NMR (125 MHz, DMSO) δ 173.29, 170.53, 169.17, 168.71, 167.79, 164.76, 160.80, 159.77, 156.81,

146.30, 143.18, 139.80, 139.62, 136.65, 135.89, 132.53, 129.87, 126.57, 124.31, 123.72, 121.92, 121.67, 121.39, 120.54, 119.65, 119.62, 117.89, 111.39, 110.32, 100.46, 49.03, 45.62, 40.59, 39.00, 31.45, 29.30, 29.26, 24.32, 22.63. LC/MS (ESI) m/z calculated $[M+H]^+$ 872.25, found 872.72.

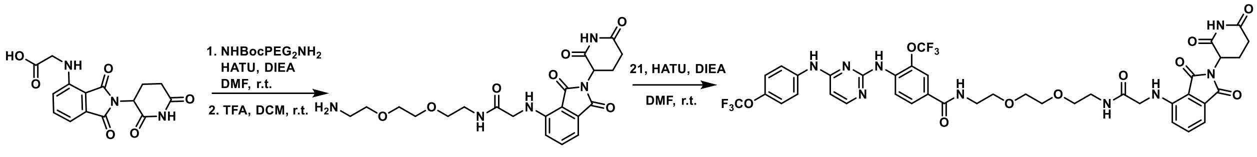


**Synthesis of *N*-(2-(2-(2-(2-((2-(2,6-dioxopiperidin-3-yl)-1,3-dioxoisoindolin-4-yl)amino)acetamido)ethoxy)ethoxy)ethyl)-3-(trifluoromethoxy)-4-((4-((4-(trifluoromethoxy)phenyl)amino)pyrimidin-2-yl)amino)benzamide (ZNL-07-054)**

Compound **ZNL-07-054** was synthesized by starting with (2-(2,6-dioxopiperidin-3-yl)-1,3-dioxoisoindolin-4-yl)glycine (**22**) and linker *tert*-butyl (2-(2-(2-aminoethoxy)ethoxy)ethyl)carbamate following the same procedure as **ZNL-07-056** was prepared as a yellow solid. $^1H$ NMR (500 MHz, DMSO-$d_6$) δ 11.10 (s, 1H), 9.65 (s, 1H), 8.81 (s, 1H), 8.63 (t, J = 5.6 Hz, 1H), 8.20 – 8.12 (m, 2H), 8.07 (d, J = 5.7 Hz, 1H), 7.86 (d, J = 7.8 Hz, 2H), 7.78 – 7.72 (m, 2H), 7.58 (dd, J = 8.5, 7.1 Hz, 1H), 7.25 (d, J = 8.6 Hz, 2H), 7.07 (d, J = 7.1 Hz, 1H), 6.95 (t, J = 5.7 Hz, 1H), 6.86 (d, J = 8.6 Hz, 1H), 6.32 (d, J = 5.8 Hz, 1H), 5.08 (dd, J = 12.8, 5.4 Hz, 1H), 3.94 (d, J = 5.7 Hz, 2H), 3.59 – 3.49 (m, 6H), 3.44 (dt, J = 7.3, 5.6 Hz, 4H), 3.26 (q, J = 5.7 Hz, 2H), 2.95 – 2.84 (m, 1H), 2.64 – 2.53 (m, 2H), 2.08 –1.98 (m, 1H). $^{13}C$ NMR (125 MHz, DMSO) δ 173.28, 170.52, 169.17, 169.07, 167.78, 165.02, 160.82, 159.70, 156.79, 146.28, 143.20, 139.61, 136.65, 136.01, 132.51, 129.44, 126.63, 124.09, 123.72, 121.93, 121.68, 121.41, 120.58, 119.65, 119.62, 117.91, 117.57, 111.42, 110.32, 100.52, 70.05, 70.02, 69.43, 69.40, 49.03, 45.61, 40.58, 39.10, 31.45, 22.62. LC/MS (ESI) m/z calculated $[M+H]^+$ 918.26, found 918.15.

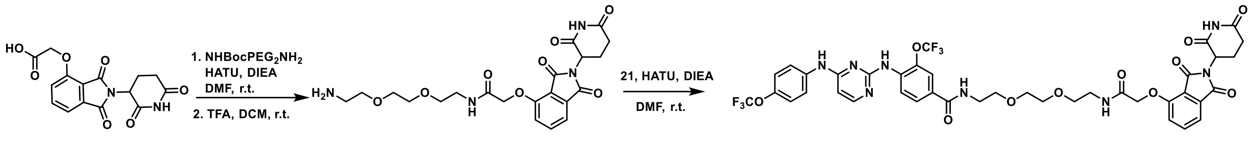


**Synthesis of *N*-(2-(2-(2-(2-((2-(2,6-dioxopiperidin-3-yl)-1,3-dioxoisoindolin-4-yl)oxy)acetamido)ethoxy)ethoxy)ethyl)-3-(trifluoromethoxy)-4-((4-((4-(trifluoromethoxy)phenyl)amino)pyrimidin-2-yl)amino)benzamide (ZNL-07-055)**

Compound **ZNL-07-055** was synthesized by starting with 2-((2-(2,6-dioxopiperidin-3-yl)-1,3-dioxoisoindolin-4-yl)oxy)acetic acid (**25**) and linker *tert*-butyl (2-(2-(2-aminoethoxy)ethoxy)ethyl)carbamate following the same procedure as **ZNL-07-056** was prepared as a yellow solid. $^1H$ NMR (500 MHz, DMSO-$d_6$) δ 11.12 (s, 1H), 9.69 (s, 1H), 8.84 (s, 1H), 8.62 (t, J = 5.6 Hz, 1H), 8.14 (d, J = 8.6 Hz, 1H), 8.07 (d, J = 5.8 Hz, 1H), 8.01 (t, J = 5.7 Hz, 1H), 7.86 (d, J = 7.8 Hz, 2H), 7.80 (dd, J = 8.5, 7.3 Hz, 1H), 7.77 – 7.73 (m, 2H), 7.49 (d, J = 7.2 Hz, 1H), 7.39 (d, J = 8.5 Hz, 1H), 7.26 (d, J = 8.6 Hz, 2H), 6.33 (d, J = 5.8 Hz, 1H), 5.12 (dd, J = 12.8, 5.4 Hz, 1H), 4.79 (s, 2H), 3.59 – 3.50 (m, 6H), 3.45 (dt, J = 14.5, 5.8 Hz, 4H), 3.31 (d, J = 5.9 Hz, 2H),

2.90 (ddd, J = 16.6, 13.7, 5.4 Hz, 1H), 2.64 – 2.53 (m, 2H), 2.09 – 2.00 (m, 1H). $^{13}$C NMR (125 MHz, DMSO) δ 173.24, 170.34, 167.37, 167.20, 165.91, 164.99, 160.84, 159.53, 156.53, 155.45, 143.26, 139.66, 139.54, 137.38, 135.90, 133.51, 129.55, 126.64, 124.15, 123.71, 121.94, 121.67, 121.47, 120.80, 120.59, 119.65, 119.62, 117.61, 117.23, 116.49, 100.53, 70.06, 69.40, 69.30, 67.96, 49.27, 38.86, 31.41, 22.46. LC/MS (ESI) m/z calculated $[M+H]^+$ 919.24, found 919.80.

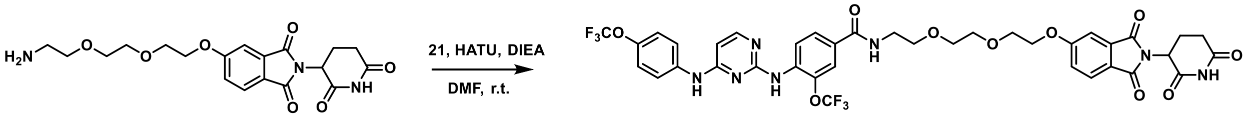


6b

ZXH-8-001

**Synthesis of *N*-(2-(2-(2-((2-(2,6-dioxopiperidin-3-yl)-1,3-dioxoisoindolin-5-yl)oxy)ethoxy)ethoxy)ethyl)-3-(trifluoromethoxy)-4-((4-((4-(trifluoromethoxy)phenyl)amino)pyrimidin-2-yl)amino)benzamide (ZXH-8-001)**

5-(2-(2-(2-aminoethoxy)ethoxy)ethoxy)-2-(2,6-dioxopiperidin-3-yl)isoindoline-1,3-dione (10mg, 0.025 mmol) and 4-((4-((4-(trifluoromethoxy)phenyl)amino)pyrimidin-2-yl)amino)-3-(trifluoromethyl)benzoic acid (**21**) (12 mg, 0.025 mmol) were added to DMF (1 mL) followed by adding DIEA (18 µL, 0.125 mmol) and HATU (19 mg, 0.05 mmol), the reaction stirred for 30 minutes and then purified by HPLC to provide the title compound **ZXH-8-001** (8 mg, 39%). $^{1}$H NMR (500 MHz, DMSO-$d_6$) δ 11.11 (s, 1H), 9.65 (s, 1H), 8.79 (s, 1H), 8.63 (t, J = 5.6 Hz, 1H), 8.14 (d, J = 8.9 Hz, 1H), 8.06 (d, J = 5.8 Hz, 1H), 7.85 (d, J = 7.7 Hz, 2H), 7.80 (d, J = 8.3 Hz, 1H), 7.77 – 7.70 (m, 2H), 7.42 (d, J = 2.3 Hz, 1H), 7.32 (dd, J = 8.3, 2.3 Hz, 1H), 7.25 (d, J = 8.2 Hz, 1H), 6.32 (d, J = 5.8 Hz, 1H), 5.11 (dd, J = 12.8, 5.4 Hz, 1H), 4.28 – 4.25 (m, 1H), 3.81 – 3.75 (m, 2H), 3.67 – 3.58 (m, 4H), 3.56 (t, J = 6.0 Hz, 2H), 3.47 – 3.43 (m, 2H), 2.88 (ddd, J = 16.9, 13.9, 5.5 Hz, 1H), 2.63 – 2.52 (m, 2H), 2.04 (ddd, J = 10.6, 5.7, 3.3 Hz, 1H). $^{13}$C NMR (125 MHz, DMSO) δ 173.26, 170.41, 167.31, 167.26, 165.02, 164.36, 160.80, 159.68, 156.80, 143.21, 139.57, 135.98, 134.36, 129.44, 126.62, 125.71, 124.06, 123.51, 121.93, 121.67, 121.43, 121.29, 120.57, 119.64, 109.31, 100.51, 70.33, 70.10, 69.37, 69.09, 68.85, 49.42, 31.40, 22.52. LC/MS (ESI) m/z calculated $[M+H]^+$ 862.22, found 862.73.

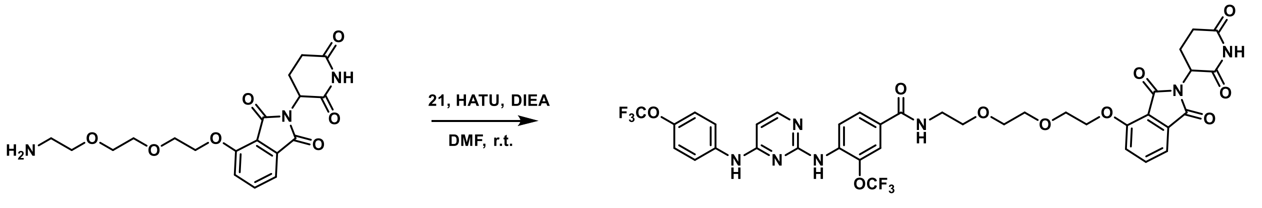


9b

ZXH-8-004

**Synthesis of *N*-(2-(2-(2-((2-(2,6-dioxopiperidin-3-yl)-1,3-dioxoisoindolin-4-yl)oxy)ethoxy)ethoxy)ethyl)-3-(trifluoromethoxy)-4-((4-((4-(trifluoromethoxy)phenyl)amino)pyrimidin-2-yl)amino)benzamide (ZXH-8-004)**

4-(2-(2-(2-aminoethoxy)ethoxy)ethoxy)-2-(2,6-dioxopiperidin-3-yl)isoindoline-1,3-dione (27mg, 0.067 mmol) and 4-((4-((4-(trifluoromethoxy)phenyl)amino)pyrimidin-2-yl)amino)-3-(trifluoromethyl)benzoic acid (**21**) (32 mg, 0.067 mmol) were added to DMF (1 mL) followed by adding DIEA (60 µL, 0.34 mmol) and HATU (51 mg, 0.13 mmol), the reaction stirred for 30 minutes and then purified by HPLC to provide the title compound **ZXH-8-004** (30 mg, 52%). $^{1}$H

NMR (500 MHz, DMSO-$d_6$) δ 11.10 (s, 1H), 10.08 (s, 1H), 9.31 (s, 1H), 8.68 (t, J = 5.6 Hz, 1H), 8.10 – 8.05 (m, 2H), 7.92 – 7.86 (m, 2H), 7.79 (dd, J = 8.5, 7.3 Hz, 1H), 7.74 – 7.67 (m, 2H), 7.50 (d, J = 8.5 Hz, 1H), 7.45 (d, J = 7.2 Hz, 1H), 7.27 (d, J = 8.1 Hz, 1H), 6.39 (d, J = 6.2 Hz, 1H), 5.08 (dd, J = 12.8, 5.4 Hz, 1H), 4.35 – 4.29 (m, 1H), 3.83 – 3.78 (m, 2H), 3.67 (dd, J = 5.9, 3.6 Hz, 2H), 3.61 – 3.53 (m, 4H), 3.44 (q, J = 5.8 Hz, 2H), 2.88 (ddd, J = 16.8, 13.8, 5.4 Hz, 1H), 2.63 – 2.52 (m, 2H), 2.50 – 2.47 (m, 2H), 2.06 – 1.99 (m, 1H). $^{13}$C NMR (125 MHz, DMSO) δ 173.25, 170.41, 167.27, 165.74, 164.83, 161.03, 158.56, 158.30, 156.28, 143.89, 140.32, 138.73, 137.42, 133.71, 130.77, 126.79, 125.11, 123.66, 122.18, 121.97, 121.63, 121.61, 120.68, 120.44, 119.60, 119.55, 116.78, 115.86, 100.62, 70.56, 70.18, 69.37, 69.30, 69.16, 49.22, 31.41, 22.45. LC/MS (ESI) m/z calculated $[M+H]^+$ 862.22, found 862.38.

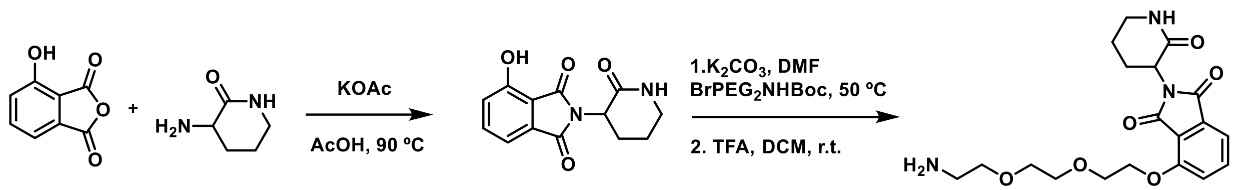


30 15 31 32

**Synthesis of 4-(2-(2-(2-aminoethoxy)ethoxy)ethoxy)-2-(2-oxopiperidin-3-yl)isoindoline-1,3-dione (32)**

A mixture of 4-hydroxyisobenzofuran-1,3-dione (1 g, 6.1 mmol), 3-aminopiperidin-2-one hydrochloride (1 g, 6.7 mmol), and NaOAc (1.8 g, 18.6 mmol) was dissolved in HOAc (20 mL), and the resulting mixture was stirred at 90 °C for 16 h. After the reaction was completed, the solution was filtered and the filtration residue was washed by water and dried to yield a yellow solid (**31**), which was used in the next step without further purification (950 mg, 60%).

4-hydroxy-2-(2-oxopiperidin-3-yl)isoindoline-1,3-dione (50mg, 0.17 mmol) was dissolved in DMF (1.5 mL), treated with $K_2CO_3$ (46 mg, 0.34 mmol), *tert*-butyl (2-(2-(2-bromoethoxy)ethoxy)ethyl)carbamate (55 mg, 0.17 mmol) was added, and the mixture stirred 6 hours at 50 ºC. The reaction was diluted with water and extracted with EtOAc. Combined extracts were washed with brine, dried with $Na_2SO_4$, then concentrated and purified by silica gel chromatography to obtain *tert*-butyl (2-(2-(2-((1,3-dioxo-2-(2-oxopiperidin-3-yl)isoindolin-4-yl)oxy)ethoxy)ethoxy)ethyl)carbamate (29 mg, 35%). LC/MS (ESI) m/z calculated $[M+H]^+$ 492.23, found [M+H-Boc] 392.45.

*tert*-butyl (2-(2-(2-((1,3-dioxo-2-(2-oxopiperidin-3-yl)isoindolin-4-yl)oxy)ethoxy)ethoxy)ethyl)carbamate (29mg, 0.06 mmol) was dissolved in 2 mL of DCM/TFA (1:1) and stirred for 2 hours at room temperature before being concentrated and dried to provide 4-(2-(2-(2-aminoethoxy)ethoxy)ethoxy)-2-(2-oxopiperidin-3-yl)isoindoline-1,3-dione(**32**). LC/MS (ESI) m/z calculated $[M+H]^+$ 392.17, found 392.45.

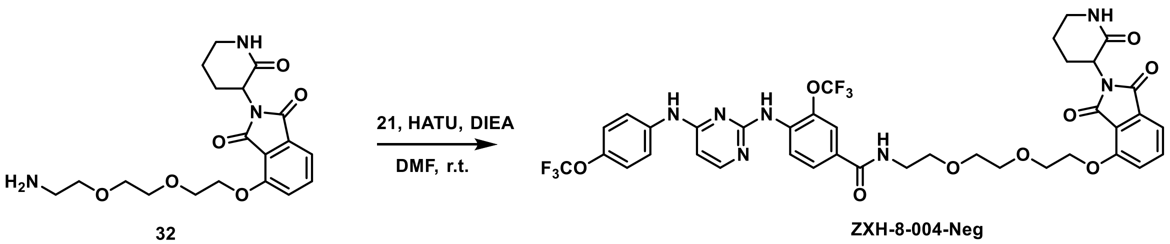


**Synthesis of *N*-(2-(2-(2-((1,3-dioxo-2-(2-oxopiperidin-3-yl)isoindolin-4-yl)oxy)ethoxy)ethoxy)ethyl)-3-(trifluoromethoxy)-4-((4-((4-(trifluoromethoxy)phenyl)amino)pyrimidin-2-yl)amino)benzamide (ZXH-8-004-Neg)**

4-(2-(2-(2-aminoethoxy)ethoxy)ethoxy)-2-(2-oxopiperidin-3-yl)isoindoline-1,3-dione (25mg, 0.064 mmol) and 4-((4-((4-(trifluoromethoxy)phenyl)amino)pyrimidin-2-yl)amino)-3-(trifluoromethyl)benzoic acid (**21**) (32 mg, 0.064 mmol) (29 mg, 0.064 mmol) were added to DMF (1 mL) followed by adding DIEA (55 µL, 0.32 mmol) and HATU (50 mg, 0.128 mmol), the reaction stirred for 30 minutes and then purified by HPLC to provide the title compound (26 mg, 48%). $^{1}$H NMR (500 MHz, DMSO-$d_6$) δ 10.60 (s, 1H), 9.96 (s, 1H), 8.74 (t, J = 5.6 Hz, 1H), 8.08 (d, J = 6.7 Hz, 1H), 7.99 (d, J = 8.4 Hz, 1H), 7.93 (dd, J = 11.7, 3.2 Hz, 2H), 7.83 (d, J = 3.0 Hz, 1H), 7.76 (dd, J = 8.5, 7.3 Hz, 1H), 7.64 (d, J = 8.7 Hz, 2H), 7.47 (d, J = 8.5 Hz, 1H), 7.41 (d, J = 7.2 Hz, 1H), 7.28 (d, J = 8.6 Hz, 2H), 6.46 (d, J = 6.7 Hz, 1H), 4.54 (dd, J = 12.0, 6.3 Hz, 1H), 4.30 (t, J = 4.5 Hz, 2H), 3.82 – 3.77 (m, 2H), 3.67 (dd, J = 5.9, 3.6 Hz, 2H), 3.61 – 3.54 (m, 4H), 3.45 (q, J = 5.8 Hz, 2H), 3.21 (dt, J = 10.1, 5.0 Hz, 2H), 2.20 (qd, J = 12.2, 4.2 Hz, 1H), 2.03 – 1.93 (m, 1H), 1.91 – 1.84 (m, 2H). $^{13}$C NMR (125 MHz, DMSO) δ 167.51, 166.02, 164.65, 161.31, 158.85, 158.57, 158.30, 156.14, 144.77, 141.24, 137.63, 137.17, 133.92, 133.24, 132.47, 126.99, 126.46, 123.16, 122.00, 121.58, 121.53, 120.81, 120.26, 119.54, 119.48, 116.96, 115.66, 100.76, 70.54, 70.18, 69.34, 69.25, 69.18, 49.27, 41.86, 40.58, 26.18, 22.17. LC/MS (ESI) m/z calculated $[M+H]^+$ 848.24, found 848.93.

**Note**

In our study, we have investigated the compounds known by various names in the follow up study. To ensure clarity, we provide here a detailed mapping of the different nomenclatures used to refer to these compounds.

| Compound Names | Other Synonyms |
|---|---|
| ZXH-2-107 | GNF-2-deg |
| ZXH-2-107-Neg | GNF-2-deg-BUMP |
| ZXH-8-004 | 2-12-2-deg |
| ZXH-8-004-Neg | 2-12-2-deg-BUMP |

**Molecular Docking**

The crystal structure of the dengue virus envelope protein bound to octyl-β-D-glucoside (βOG) (PDB code: 1OKE) was used to model the compound binding, and the hydrophobic pocket occupied by β-OG was chosen as the putative small molecule binding site. The protein was prepared with default protein preparation protocol in Schrodinger suite (2021-3 release) including adding hydrogen atoms and missing side chains, and constraint energy minimization prior to docking. Docking was carried out using Glide (Schrödinger suite), with enhanced sampling option in compound conformation and placement sampling. Ligand molecules were prepared by LigPrep with default options. Five top poses of each molecule were output for visual inspection. For each molecule, the best scored binding pose that was consistent with the binding site property, across ligands, and previously known compound SAR was chosen for further refinement by ligand-receptor refinement procedure in Schrödinger, which optimized the ligand interactions with residues within 5 Å.

**Cell culture and virus**

All the mammalian cell lines are cultured in Dulbecco's Modified Eagle's medium (DMEM) supplemented with nonessential amino acids and 10% fetal bovine serum (FBS) and incubated in a 37 °C incubator with 5% $CO_2$. Huh7.5 cells are obtained from Charles Rice (Rockefeller University) and Baby Hamster Kidney cells (BHK) are obtained from Eva Harris (University of California, Berkeley). C6/36 cells, a mosquito cell line derived from Aedes albopictus (Diptera: Culicidae) embryonic tissue (ATCC), are maintained in Leibovitz medium (L-15) containing 10% FBS at 28 °C. All the works related to infectious viruses are performed under a biosafety level 2 laboratory (BSL2). Dengue virus serotype 2 New Guinea C (DENV2 NCG) is obtained from Lee Gehrke (Massachusetts Institute of Technology) and propagated in C6/36 cells as previously described.[1, 2]

**Antibodies**

Monoclonal antibody 4G2 targeting the conserved fusion loop of the E protein was collected from culture media of hybridoma D1-4G2-4-15 (ATCC HB-112). Mouse monoclonal antibody anti-

GAPDH was purchased from GeneTex (GTX28245) and used at a ratio of 1:10000. Rabbit polyclonal anti-Abl antibody was purchased from Cell Signaling Technology (2862S) and used at a ratio of 1:1000. Horseradish peroxidase (HRP)-conjugated goat anti-mouse IgG (170-6516) and anti-rabbit IgG antibodies (170-6515) were purchased from Bio-Rad Laboratories and used at a ratio of 1:3000.

**Cell viability assays**

Cell proliferation assays were conducted as previously described.[3] K562 cells were seeded at the density of 800 cells/well in 384 well plate (corning, #3570). Then drugs were added at indicated concentrations by the HP D300 digital dispenser (Tecan). After treatment for three days, cell viability was measured using the CellTiter-Glo® luminescent cell viability assay kit (Promega, #G7570) according to the manufacturer's instructions.

**Antiviral activity**

50,000 Huh7.5 cells were seeded into each well of a 24 well plate (Corning, #3524) and incubated for 24h. The cells were then infected with each of the four DENV serotypes, DENV1 WP74, DENV2 NGC, DENV3 THD3, or DENV4 TVP360 at an MOI 0.5 or 1 for 1h, and then washed with PBS to remove extracellular virus inoculum. The infected cells were then treated with compounds at 5 and 2.5 μM concentrations in Dulbecco's Modified Eagle's medium (DMEM, Corning #10-013-CV) supplemented with nonessential amino acids, 10 mM HEPES, and 2% fetal bovine serum (FBS). At 24h post-infection, the cell culture supernatants were collected. Viral titers of supernatants were quantified by plaque formation assay (PFA). [2, 4] BHK21 cells, cultured in Minimum Essential Medium-Alpha (MEM-α, Corning #10-022-CV) with 5% FBS, were used in the PFA to determine titers of DENV1 WP74, DENV2 NGC, and DENV4 TVP360. Vero cells cultured in DMEM with 2% FBS were used to titer DENV3 THD3.

**CRBN cellular engagement assay**

HEK293T cells stably expressing the $BRD4_{BD2}$-GFP with mCherry reporter were seeded in 384-well plates (Corning REF 3764) at 5000 cells per well in 50μM. FluoroBrite DMEM media (Thermo Fisher Scientific A18967) containing 10% FBS a day before compound treatment. The cells were pretreated with the compounds via a D300e Digital Dispenser (HP), normalized to 0.5% DMSO, and incubated with the cells for 1 hour. After 1hour incubation, the cells were treated with 100nM dBET6 using the D300e Digital Dispenser and incubated together for an additional 5 hours. The assay plate was imaged immediately following the 6-hour total incubation using an ImageXpress confocal microscope with a 10X objective lens with 488 nm and 561 nm lasers in a 2uM x 1μM grid per well format. The resulting images were analyzed using a series of image analysis steps. First, the red and green channels were aligned and cropped to target the middle of each well (to avoid analysis of heavily clumped cells at the edges), and a background illumination function was calculated for both red and green channels of each well individually and subtracted to correct for illumination variations across the 384-well plate from various sources of error. An additional step was then applied to the green channel to suppress the analysis of large auto fluorescent artifacts and enhance the analysis of cell specific fluorescence by way of selecting for objects under a given size, 30 A.U., and with a given shape, speckles. mCherry-positive cells were then identified in the red channel filtering for objects between 8-60 pixels in diameter and using intensity to distinguish between clumped objects. The green channel was then segmented into GFP positive and negative areas and objects were labeled as GFP positive if at least 40% of it overlapped with a GFP positive area. The fraction of GFP-positive cells/mCherry-positive cells in each well was then calculated. The values for the concentrations that lead to a 50% increase in $BRD4_{BD2}$-eGFP accumulation ($EC_{50}$) were calculated using the nonlinear fit variable slope model (GraphPad Software).[5]

**Cellular GSPT1 degradation assay**

Cellular GSPT1 degradation assays were conducted as previously described.[6] Stable cells expressing the GSPT1-eGFP protein fusion and the mCherry reporter were seeded at a density of 30,000 cells/well in a 96-well plate. GSPT1-eGFP reporter cells were treated with increasing concentration of CC885 or indicated compounds for 24 hrs. GSPT1-eGFP reporter cell suspension was transferred to a 96 well round bottom plate compatible with the flow cytometer (guava easyCyte HT, Millipore). Green fluorescent signal and red florescent signal were individually measured and exported for analysis. Data were analyzed using Flowjo (FlowJo, LCC), and the values for the concentrations resulting in 50% degradation (DC50) were calculated using the nonlinear fit variable slope model (GraphPad Software). Data are represented as means ± s.d of duplicates (n = 2).

**Cellular thermal shift assay (CETSA)**

Huh7.5 cells were seeded at a density of 1,000,000 cells/well in 6 well plates. The cells were incubated for 24h at 37 ºC under 5% $CO_2$. The cells were then infected with dengue virus serotype 2 strain New Guinea C (NGC) at MOI 1 for 1h, then washed to remove extracellular virus, and overlaid with medium. The infected cells were then harvested at 24h post-infection. 1mL 1,000,000 infected cells are incubated with 10μM of each compound under 37 ºC for 1h. Following this, 100μL (or 100,000-infected cells) were aliquoted into 96 PCR well plates and heated up to variable temperature for 3mins. The cell lysates were then collected by three freeze-thaw cycles followed by centrifugation at 12,500g at 4ºC for 2mins. The abundance of E protein was then characterized by immunoblot analysis.[7]

**Proteomics**

MOLT4 cells were treated with DMSO or 3 µM ZXH-08-004 for 5 h. Cells were harvested by centrifugation and washed with phosphate buffered saline (PBS) before snap freezing in liquid nitrogen. Cells were lysed by addition of lysis buffer (8 M Urea, 50 mM NaCl, 50 mM 4-(2-hydroxyethyl)-1-piperazineethanesulfonic acid (EPPS) pH 8.5, Protease and Phosphatase inhibitors) and homogenization by bead beating (BioSpec) for three repeats of 30 seconds at 2400. Bradford assay was used to determine the final protein concentration in the clarified cell lysate. 50 µg of protein for each sample was reduced, alkylated and precipitated using methanol/chloroform as previously described [8] and the resulting washed precipitated protein was allowed to air dry. Precipitated protein was resuspended in 4 M Urea, 50 mM HEPES pH 7.4, followed by dilution to 1 M urea with the addition of 200 mM EPPS, pH 8. Proteins were first digested with LysC (1:50; enzyme:protein) for 12 h at RT. The LysC digestion was diluted to 0.5 M Urea with 200 mM EPPS pH 8 followed by digestion with trypsin (1:50; enzyme:protein) for 6 h at 37 °C. Sample digests were acidified with formic acid to a pH of 2-3 prior to desalting using C18 solid phase extraction plates (SOLA, Thermo Fisher Scientific). Desalted peptides were dried in a vacuum-centrifuged and reconstituted in 0.1% formic acid for LC-MS analysis.

Data were collected using a TimsTOF Pro2 (Bruker Daltonics, Bremen, Germany) coupled to a nanoElute LC pump (Bruker Daltonics, Bremen, Germany) via a CaptiveSpray nano-electrospray source. Peptides were separated on a reversed-phase $C_{18}$ column (25 cm x 75 µm ID, 1.6 µM, IonOpticks, Australia) containing an integrated captive spray emitter. Peptides were separated using a 50 min gradient of 2 - 30% buffer B (acetonitrile in 0.1% formic acid) with a flow rate of 250 nL/min and column temperature maintained at 50 ºC.

DDA was performed in Parallel Accumulation-Serial Fragmentation (PASEF) mode to determine effective ion mobility windows for downstream diaPASEF data collection [9]. The ddaPASEF parameters included: 100% duty cycle using accumulation and ramp times of 50 ms each, 1 TIMS-MS scan and 10 PASEF ramps per acquisition cycle. The TIMS-MS survey scan was acquired between 100 – 1700 *m/z* and 1/k0 of 0.7 - 1.3 V.s/cm$^2$. Precursors with 1 – 5 charges were selected and those that reached an intensity threshold of 20,000 arbitrary units were actively excluded for 0.4 min. The quadrupole isolation width was set to 2 *m/z* for *m/z* <700 and 3 *m/z* for *m/z* >800, with the *m/z* between 700-800 *m/z* being interpolated linearly. The TIMS elution voltages were calibrated linearly with three points (Agilent ESI-L Tuning Mix Ions; 622, 922, 1,222 *m/z*) to determine the reduced ion mobility coefficients ($1/K_0$). To perform diaPASEF, the precursor distribution in the DDA *m/z*-ion mobility plane was used to design an acquisition scheme for DIA data collection which included two windows in each 50 ms diaPASEF scan. Data was acquired using sixteen of these 25 Da precursor double window scans (creating 32 windows) which covered the diagonal scan line for doubly and triply charged precursors, with singly charged precursors able to be excluded by their position in the m/z-ion mobility plane. These precursor isolation windows were defined between 400 - 1200 *m/z* and 1/k0 of 0.7 - 1.3 V.s/cm$^2$.

**LC-MS data analysis**

The diaPASEF raw file processing and controlling peptide and protein level false discovery rates, assembling proteins from peptides, and protein quantification from peptides was performed using library free analysis in DIA-NN 1.8 [10]. Library free mode performs an in silico digestion of a given protein sequence database alongside deep learning-based predictions to extract the DIA precursor data into a collection of MS2 spectra. The search results are then used to generate a spectral library which is then employed for the targeted analysis of the DIA data searched against a Swissprot human database (January 2021). Database search criteria largely followed the default settings for directDIA including: tryptic with two missed cleavages, carbomidomethylation of cysteine, and oxidation of methionine and precursor Q-value (FDR) cut-off of 0.01. Precursor quantification strategy was set to Robust LC (high accuracy) with RT-dependent cross run normalization. Proteins with missing values in any of the treatments and with poor quality data were excluded from further analysis (summed abundance across channels of <100 and mean number of precursors used for quantification <2). Protein abundances were scaled using in-house scripts in the R framework [11] and statistical analysis was carried out using the limma package within the R framework [12].

**Supplementary figures**

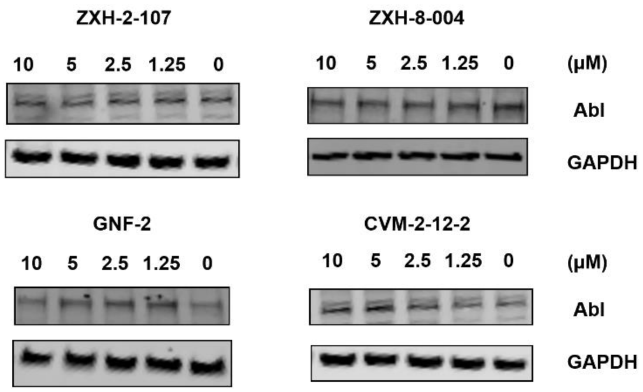


**Figure S1.** Immunoblot analysis of ABL in Huh 7.5 cells treated with ZXH-2-107, ZXH-8-004, GNF-2, and CVM-2-12-2 for 8 h.

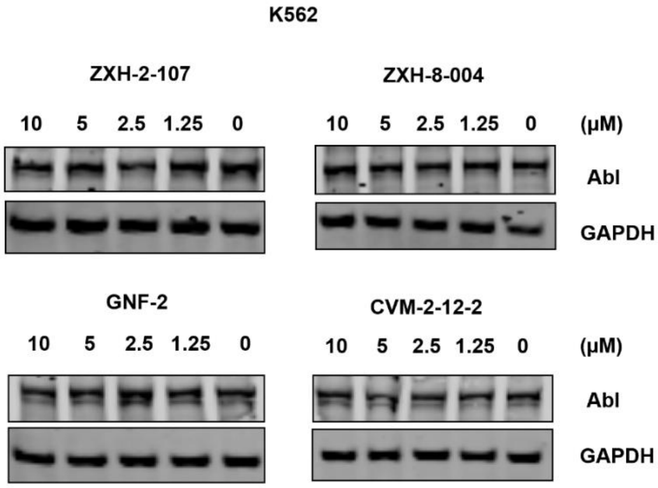


**Figure S2.** Immunoblot analysis of ABL in K562 cells treated with GNF-2, CVM-2-12-2, ZXH-2-107, ZXH-2-107-Neg, ZXH-8-004, and ZXH-8-004-Neg at indicated concentrations for 8 h.

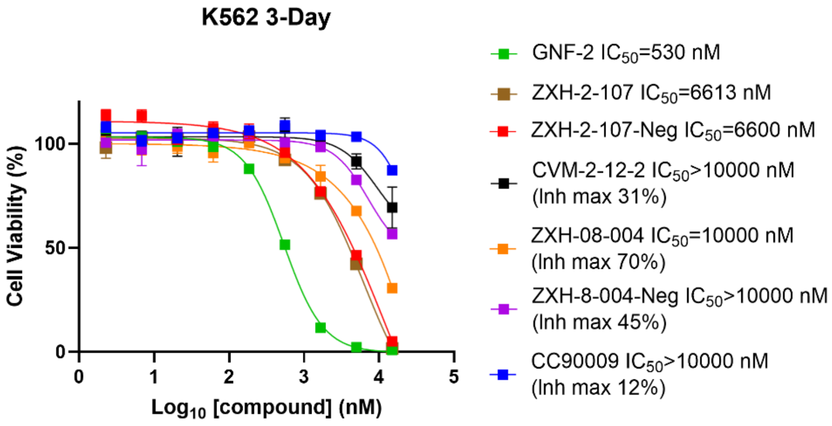


**Figure S3.** Anti-proliferation assay examining the growth inhibitory effect of a dose escalation of GNF-2, ZXH-2-107, ZXH-2-107-Neg, CVM-2-12-2, ZXH-8-004, ZXH-8-004-Neg, and CC-90009 against K562 cells for 72 hours.

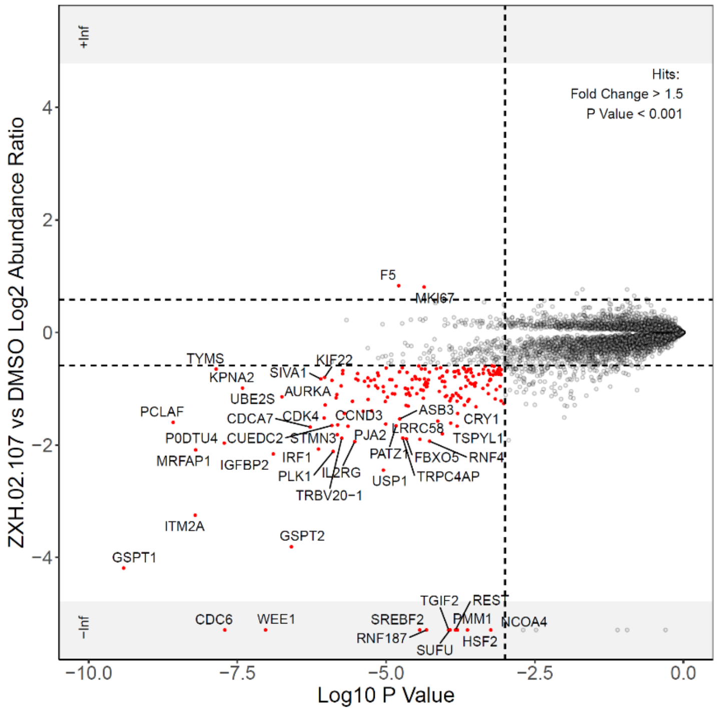


**Figure S4**. Proteome-wide degradation selectivity of ZXH-2-107 at a dose of 3 µM in MOLT4 cells after 5 hours treatment.

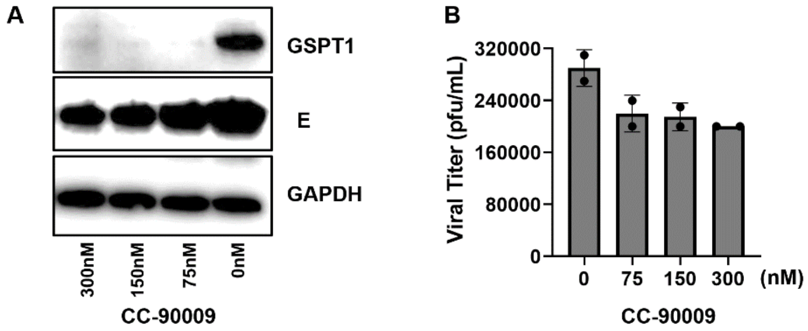


**Figure S5**. A) Immunoblot analysis of E protein in DENV2-infected Huh 7.5 cells treated with CC-90009 with different doses for 24 h. B) Antiviral activities of CC-90009 in DENV2 infected Huh 7.5 cells at different doses.

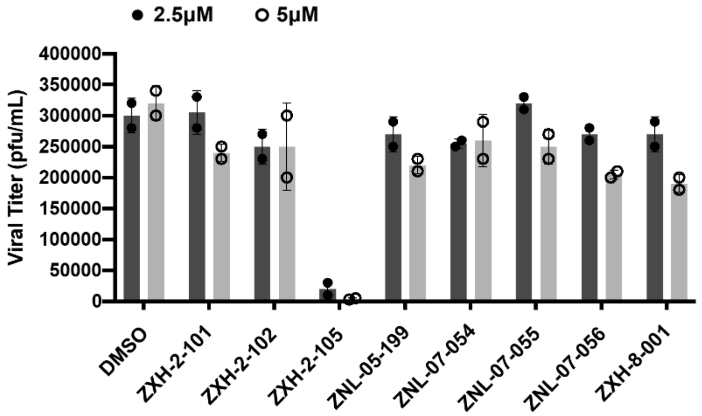


**Figure S6.** Antiviral activities of all the compounds in DENV2 infected Huh 7.5 cells at 2.5 µM and 5 µM.

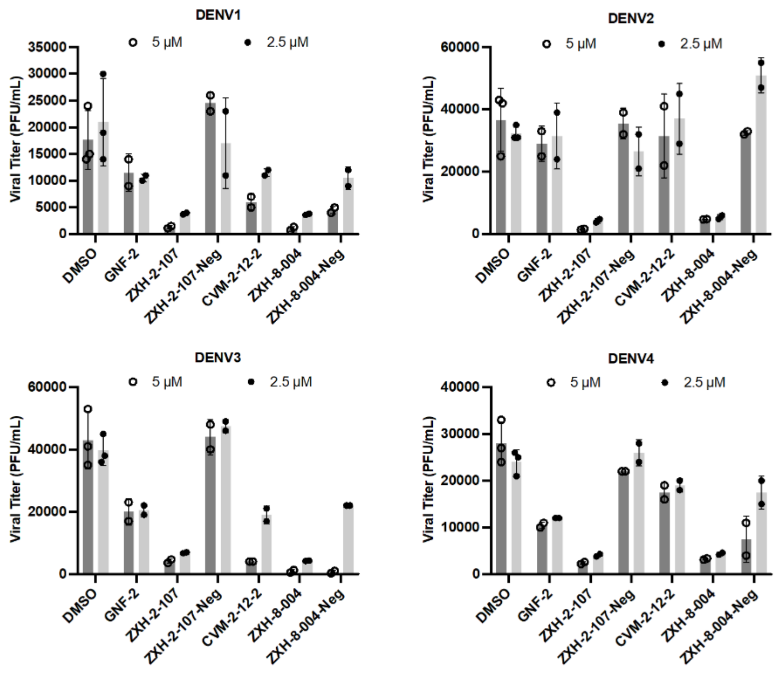


**Figure S7.** Antiviral activities of E degraders, negative controls, and parental inhibitors in DENV1, 2, 3, and 4 infected Huh 7.5 cells at 2.5 μM and 5 μM. Graphs show Viral titer in PFU/mL in Y-axis.

**Table S1.** MDR1-MDCK II permeability of lead E protein degraders and parental compounds.

| Compound | MDR1-MDCK II A2B[a] | MDR1-MDCK II B2A[a] | MDR1-MDCK II ER[b] |
|---|---|---|---|
| GNF-2 | 6.08 | 2.86 | 0.47 |
| CVM-2-12-2 | <0.216 | <0.194 | NA |
| ZXH-2-107 | 0.189 | 9.92 | 52.5 |
| ZXH-8-004 | 0.0696 | 2.64 | 38.0 |

[a]Permeability/$10^{-6}$ cm $s^{-1}$. [b]Efflux ratio. Note: The signal responses of CVM-2-12-2 in the receiver samples were undetectable, the accurate value of ER value cannot be calculated.
